# Large-scale analysis of post-translational modifications in E. coli under glucose-limiting conditions

**DOI:** 10.1101/051185

**Authors:** Colin W Brown, Viswanadham Sridhara, Daniel R Boutz, Maria D Person, Edward M Marcotte, Jeffrey E Barrick, Claus O Wilke

**Affiliations:** Institute for Cellular and Molecular Biology, The University of Texas at Austin, Speedway, Austin, USA.; Center for Computational Biology and Bioinformatics, The University of Texas at Austin, Speedway, Austin, USA.; Center for Systems and Synthetic Biology, The University of Texas at Austin, Speedway, Austin, USA.; College of Pharmacy, The University of Texas at Austin, Speedway, Austin, USA.; Department of Molecular Biosciences, The University of Texas at Austin, Speedway, Austin, USA.; Department of Integrative Biology, The University of Texas at Austin, Speedway, Austin, USA.

**Keywords:** post-translational modification, proteomics, prokaryote

## Abstract

**Background:** Post-translational modification (PTM) of proteins is central to many cellular processes across all domains of life, but despite decades of study and a wealth of genomic and proteomic data the biological function of many PTMs remains unknown. This is especially true for prokaryotic PTM systems, many of which have only recently been recognized and studied in depth. It is increasingly apparent that a deep sampling of abundance across a wide range of environmental stresses, growth conditions, and PTM types, rather than simply cataloging targets for a handful of modifications, is critical to understanding the complex pathways that govern PTM deposition and downstream effects.

**Results:** We utilized a deeply-sampled dataset of MS/MS proteomic analysis covering 9 timepoints spanning the *Escherichia coli* growth cycle and an unbiased PTM search strategy to construct a temporal map of abundance for all PTMs within a 400 Da window of mass shifts. Using this map, we are able to identify novel targets and temporal patterns for N-terminal Nα acetylation, C-terminal glutamylation, and asparagine deamidation. Furthermore, we identify a possible relationship between N-terminal Nα acetylation and regulation of protein degradation in stationary phase, pointing to a previously unrecognized biological function for this poorly-understood PTM.

**Conclusions:** Unbiased detection of PTM in MS/MS proteomics data facilitates the discovery of novel modification types and previously unobserved dynamic changes in modification across growth timepoints.

## Background

Post-translational modification of proteins (PTM) is a ubiquitous paradigm for dynamic cellular response and information transfer across all kingdoms of life [1]. Although historically PTM has been studied in the context of discrete and tightly-regulated signal transduction systems such as eukaryotic histone proteins [2], kinase cascades [3, 4], and prokaryotic two-component systems [5], it is only relatively recently, with the development of tandem-mass-spectrometry-based proteomics, that the abundance and complexity of PTM has become apparent [6]. A surprising result from many of these investigations has been that the phylogenetic distribution of many PTMs is much wider than had been assumed. A number of PTM types previously thought to be restricted to eukaryotic and metazoan species, such as lysine acetylation [7], serine/threonine phosphorylation [8], tyrosine phosphorylation [9, 10], and ubiquitination-like protein ligation [11], are now known to be relatively common in prokaryotic proteomes as well. This realization, in combination with the recognition that PTM plays a critical role in growth and virulence of important prokaryotic pathogens (e.g. [9, 12, 13, 14, 15, 16]), has highlighted the need for a better understanding of prokaryotic PTM and particularly the need for deeper, proteome-scale analysis of prokaryotic PTMs.

In response to these needs, much progress in recent years has been made in the mapping of important PTMs across a wide range of prokaryotes [7, 17, 1]. However, the vast majority of these studies are limited by only examining a handful of easily-achieved culture conditions and timepoints, and by only examining a single PTM type in isolation. The former limitation is especially important, as batch cultures grown for short time periods in rich media, as is most common for bacterial proteomics experiments, may be a poor reflection of the high-stress, nutrient-starved conditions in which bacteria spend most of their time in the wild [18, 19]. While collecting bacterial samples directly from their natural habitat is generally infeasible for proteomics experiments given the requirements for large cell numbers and pure samples, the starvation conditions commonly encountered in a bacterium’s native habitat are thought to be largely recapitulated in long-term batch culture [18, 19]. As an exponentially-growing batch culture exhausts the readily available nutrients in the growth medium, the cells undergo a regulated transition into stasis by activating a stereotypic stress response. This response usually involves a decrease of or complete stop to cell division, steep dropoff’s in oxidative metabolism [20] and protein synthesis [21], sequestration of ribosomes [22, 23], activation of oxidative damage response systems [24, 25], and increased protease-mediated protein turnover [26]. Eventually, even this inactive state becomes unsustainable for the majority of cells in the culture, and a large-scale die-off takes place until the culture reaches an equilibrium where the remaining cells are able to survive on the nutrients liberated from their less fortunate culture-mates. This “deep stationary” phase of batch culture is poorly understood, but is characterized by a gradual loss of culturability (the **V**iable **B**ut **N**on **C**ulturable state [27]), likely related to accumulated cell damage, and a dynamic equilibrium of genetic changes as mutations advantageous for stationary phase growth (**G**rowth **A**dvantage in **S**tationary **P**hase, or GASP mutations [19]) are fixed by selection in the population. The low rate of protein synthesis and the potential importance of nonenzymatic protein-damage modifications in stationary phase makes an understanding of PTM chemistry and dynamics during this portion of the growth cycle especially important.

With a few very recent exceptions [28, 29], studies of PTMs at different growth phases in *E. coli* have been restricted to either a single modification or a handful of pre-specified modifications (e.g. [30, 31, 32, 10]). This limitation is largely due to both the relatively low abundance of the PTMs examined, necessitating the enrichment of modified peptides using PTM-specific antibodies [6] or chromatographic separations [33], and data analysis tools that are only useful for examining a small number of pre-specified PTMs. While enrichment is necessary for relatively transient modifications such as phosphorylation, particularly where a broad survey of targets rather than PTM dynamics is the experimental goal, it has a critical shortcoming in that it makes quantitative comparisons among PTM types, and perhaps more importantly between modified and unmodified copies of an individual protein, impossible. Adding to this problem is the fact that many of the most commonly-used software packages for MS/MS spectrum-peptide sequence matching (e.g. Mascot [34], Sequest [35], OMSSA [36], or TANDEM [37]) are limited by the need to create an *in silico* database of theoretical spectra using an existing peptide library; while this approach facilitates rapid searching, it makes searches involving more than a few PTM types computationally unwieldy. The spectrum of PTMs beyond a handful of well-studied examples is therefore largely unexplored.

In this work we utilize a recently developed computational tool for unrestricted analysis of PTMs in MS/MS proteomics data, MODa [38], to examine a unique proteomic dataset [39] covering 9 timepoints of the *E. coli* REL606 growth curve in minimal glucose media from early exponential growth (3 hours post-innoculation) to deep stationary phase (336h, or 2 weeks post-innoculation). MODa uses a combination of *de novo* sequence-tag matching and spectral alignment to make assigning PTM-containing spectra across a wide range of mass shifts computationally tractable, and this allows us to construct an unbiased PTM spectrum across all phases of growth for all modifications from −200 Da to +200 Da. The fine temporal resolution of our dataset then allows us to identify novel temporal trends in a number of PTMs, including N-terminal Nα acetylation, C-terminal glutamylation, and asparagine deamidation. In addition, the lack of bias or enrichment for specific PTMs allows us to track behavior of modified and unmodified proteins across the growth cycle, and to identify a potential functional relationship between N-terminal acetylation, protein oxidative damage, and stationary-phase protein degradation.

## Results

We took advantage of a previously existing LC-MS/MS proteomics dataset [39] isolated from 3 biological replicate cultures of *E. coli* B REL606 sampled across 9 timepoints, from early exponential phase (3h post-innoculation) to extended late stationary phase (336h, or 2 weeks post-innoculation). The raw spectra from this dataset were used for simultaneous spectrum-sequence matching and PTM identification using the hybrid fragment matching/spectral alignment software MODa [38]. To reduce computation time and limit the occurrence of false positives, we restricted the MODa search to single-peptide mass shifts of +/− 200 Daltons, with one PTM allowed per peptide spectral match (PSM). To further limit the occurrence of false positive matches, we used the MODa “correct match” probability [38] to calculate the false discovery rate (FDR) and construct subsets of the highest-probability PSMs with 5% and 1% FDR (hereafter referred to as FDR5 and FDR1, respectively). The samples in our analysis were treated with IAA to modify cysteines with a 57 Da carbamidomethyl group; during MODa analysis, this was treated as a static modification to cysteine (i.e. all modifications were relative to the molecular weight of Cys + 57Da). However, this results in an incorrect mass shift for any Cys PTMs that prevent carbamidomethylation (e.g. oxidation), so we added 57 Da to all Cys modifications to ensure that mass shifts for these modifications matched those for non-Cys residues.

We identified a total of 2,527,135 PSMs across all 27 samples, corresponding to a total of 32,755 peptides that occur in at least one sample; these peptides represent 3,544 individual proteins when all timepoints are considered (Table 1). FDR filtering lowers these numbers substantially, yielding 1,980,884 PSMs and 22,776 unique peptides across 2,445 proteins in the 5% FDR set, and 1,473,636 PSMs and 19,265 unique peptides across 2121 proteins in the 1% FDR set (Table 1). These filtered numbers are in agreement with previous proteomic experiments in *E. coli* [39, 40, 31], with the slightly lower number of proteins in our analysis, likely a result of the reduced sensitivity inherent in the larger search space used by MODa.

**Table 1.**
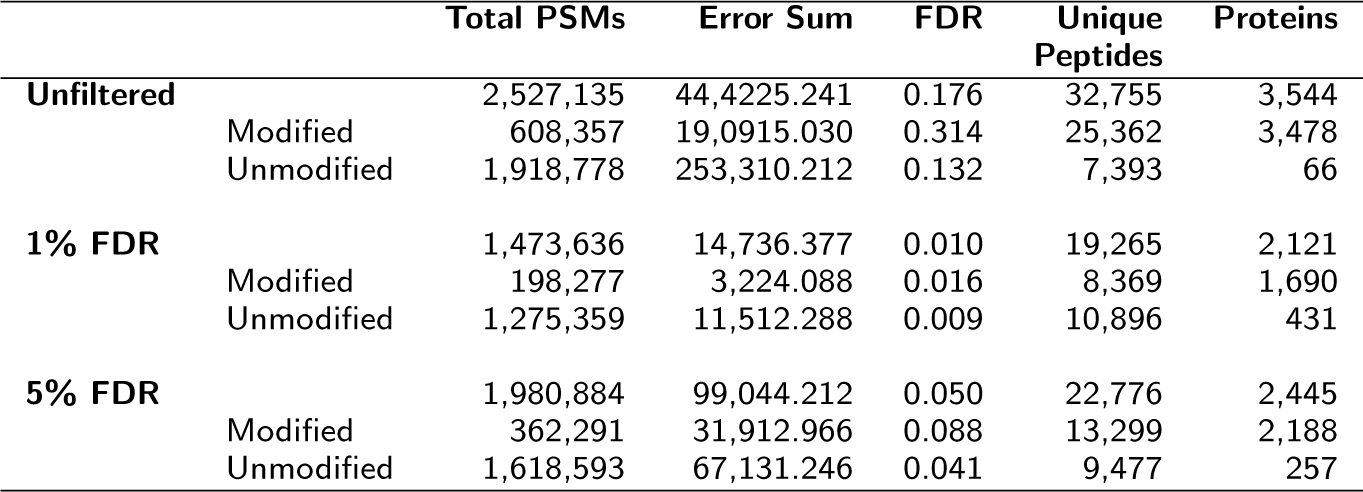
Counts of PSMs, Unique Peptides, and Proteins for Unfiltered, 1% FDR, and 5% FDR Datasets

### A large fraction of the *E. coli* proteome undergoes PTM during growth and starvation in glucose

Of the 1,473,636 PSMs identified across all timepoints in the 1% FDR dataset, a remarkably large fraction, 198,277 (13.5%), are predicted by MODa as having a putative PTM. These modified PSMs corresponded to 8,369 out of 19,265 unique peptides (42%) having at least one modification in any sample, and 1,690 out of 2121 proteins (79.7%) having at least one modification on any constituent peptide. Interestingly, the proportion of the proteome predicted to have at least one PTM remains relatively constant across time points and biological replicates. PSMs, unique peptides, and proteins all show very little change in the proportion of overall PTM across all 9 time points (Figure 1).

**Figure 1.**
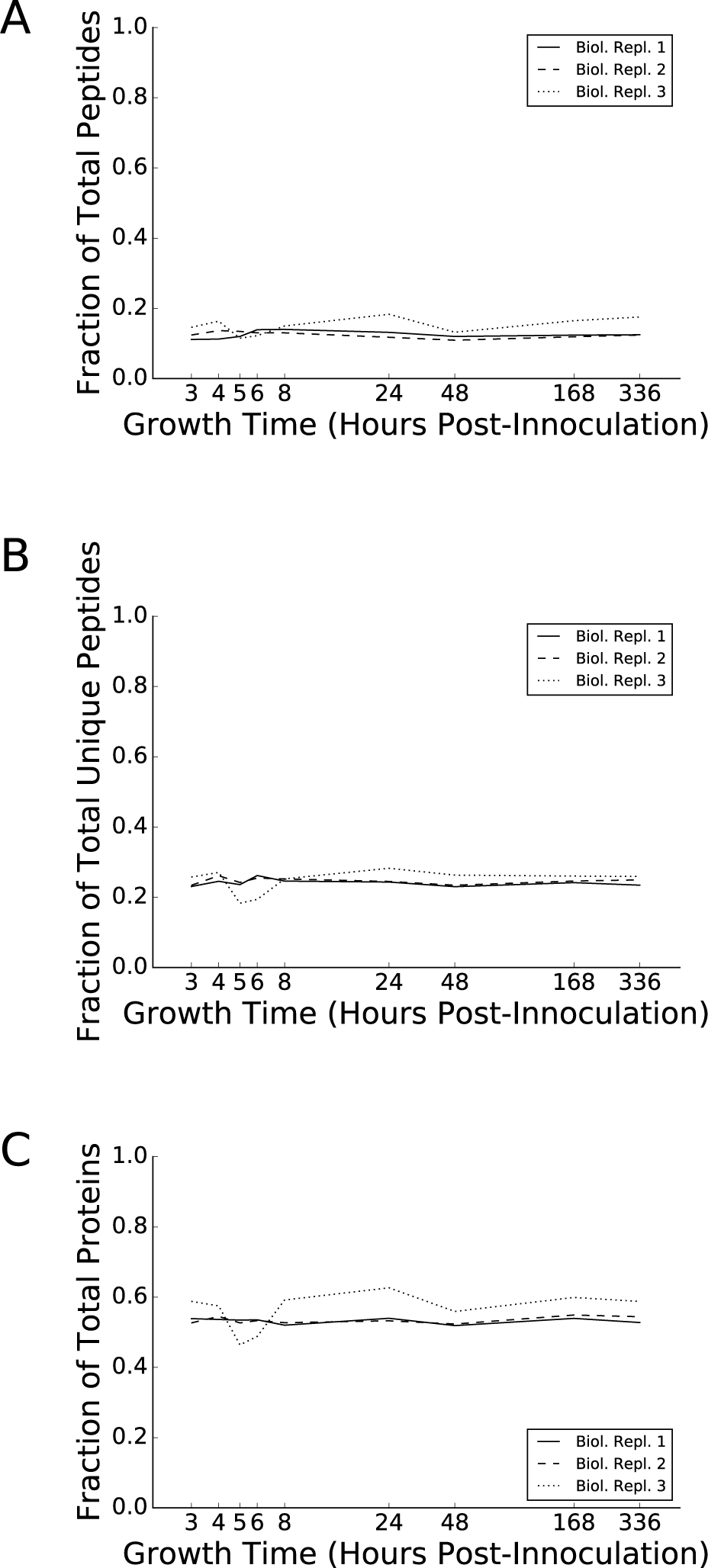
Global abundance of all modifications across growth timepoints. Fraction of total counts of PSMs (A), unique peptides (B), and proteins (C) containing at least one mass shift passing the 1% FDR threshold at the timepoint indicated on the *x*-axis for biological replicates 1, 2 and 3 (solid, dashed, and dotted lines, respectively)

### Composition of the *E. coli* PTM spectrum

A unique feature of our analysis strategy is the ability to conduct an unbiased search for spectra matching post-translationally modified peptides across a wide range of possible mass shifts. We used MODa to search our raw spectral data for 400 potential peptide mass shifts, ranging from −200 Da to +200 Da; counts of PSMs for this range of mass shifts are shown in Figures 2 and S1.

**Figure 2.**
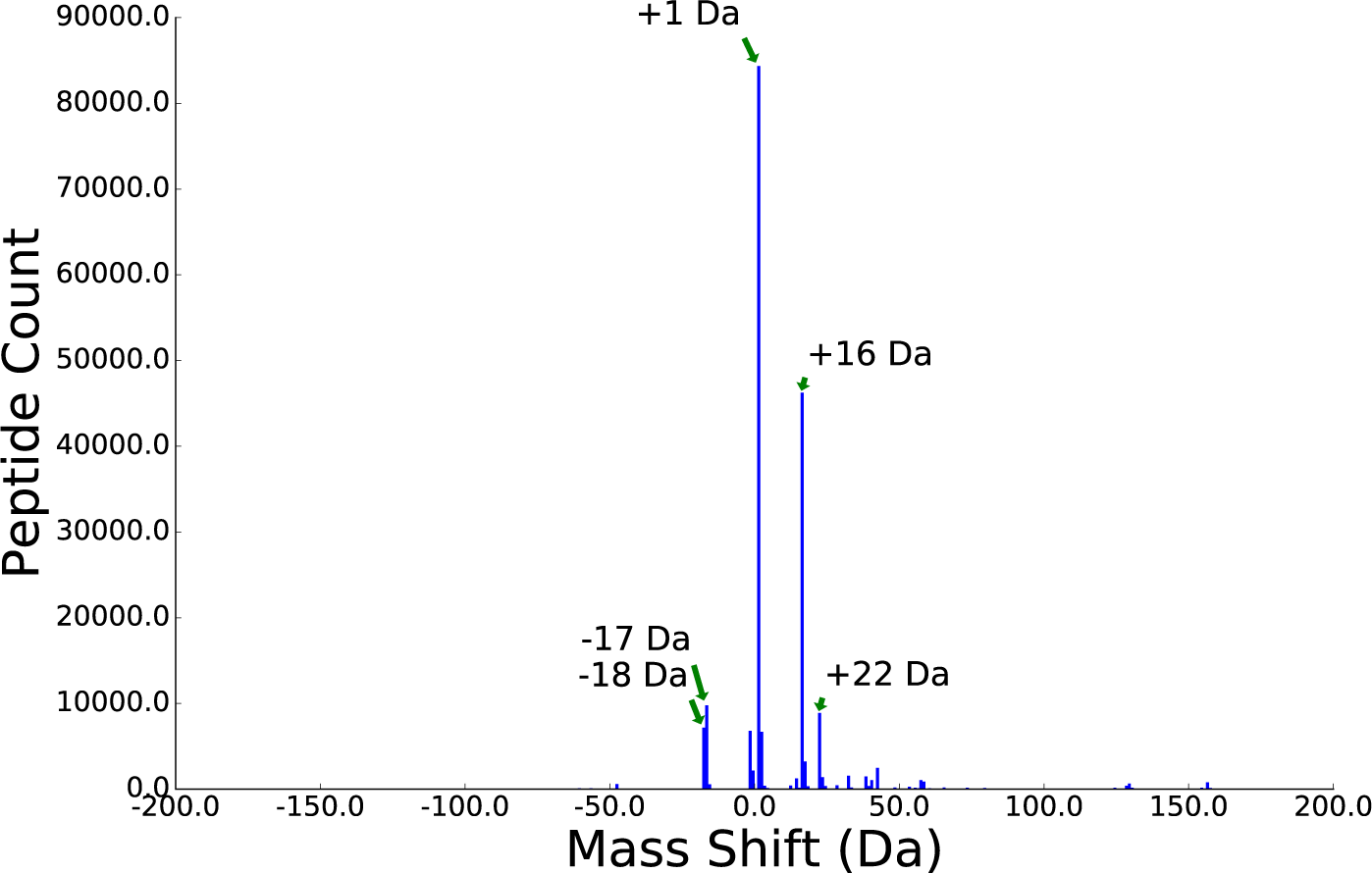
Mass-shift counts across all timepoints and datasets. PSM counts for all mass shifts identified by MODa[38] between −200 Da and +200 Da, summed across all nine timepoints and all three biological replicates; labeled peaks are the top 5 most abundant (by raw count) mass shifts in the dataset.

The overall abundance of individual mass shifts varies widely, with the most abundant mass shifts corresponding to small functional group modifications. The most abundant mass shift is a neutral gain of 1 Da (84,357 PSMs, 45% of all modified PSMs). In addition to simple hydrogenation, this mass change can result from a number of more complicated modifications and MS artifacts; see Discussion. Other abundant mass shifts include oxidations (+16 Da, 46,244 PSMs, 24% of all modified PSMs; +32 Da, 1,563, 0.8% of all modified PSMs), metal ion adducts such as sodium (+22 Da, 8,882 PSMs, 4.7% of all modified PSMs) and potassium (+38 Da, 1,490 PSMs 0.79% of all modified PSMs), and neutral losses such as deamidation (−17 Da, 9,780 PSMs, 5.2% of all modified PSMs) and dehydration (−18 Da, 7,169 PSMs, 3.8% of all modified PSMs).

Commonly studied regulatory PTMs are relatively rare in our data, most likely due to their low abundance in the proteome and the fact that our samples did not undergo enrichment for specific modifications prior to analysis. Although a large number of apparent acetylations (+42 Da) were identified, only a handful of these map to known acetylated lysine residues [32, 30, 41]. A small number of phosphorylations (+80 Da) were identified, although the majority of these are modifications to an active-site serine that acts as a phosphoryl group donor during catalysis in the metabolic enzyme phosphoglucomutase (see Table 2). A list of counts for all mass shifts recovered by MODa is included in Additional File 12.

**Table 2.**
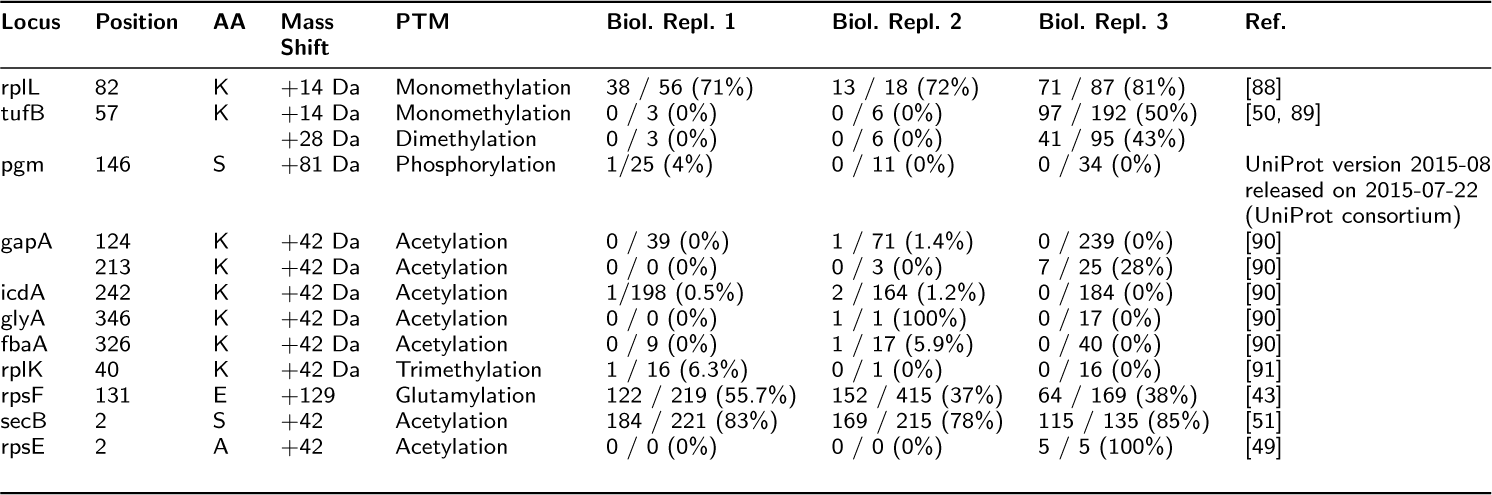
Previously Identified Post-Translational modifications recovered in our analysis

### Distribution of target amino-acid residues varies widely among mass shifts

The most commonly modified amino acid across all timepoints is methionine—nearly all of these modifications are a +16 Da shift corresponding to oxidation (see Discussion)—followed by the hydrophobic amino acids Ala, Val, Leu, Ile; amide-containing amino acids Asn and Gln; and their carboxyl counterparts Asp and Glu (Table 3). The observation of a large number of modifications on amino acids with hydrocarbon side-chains, which are generally not expected to undergo PTM, can likely be explained by a combination of incorrect assignment of a mass shift to the amino acid (AA) by MODa, modification of the backbone NH or CO groups, or selection of peaks with isotopically shifted masses during MS2.

**Table 3.**
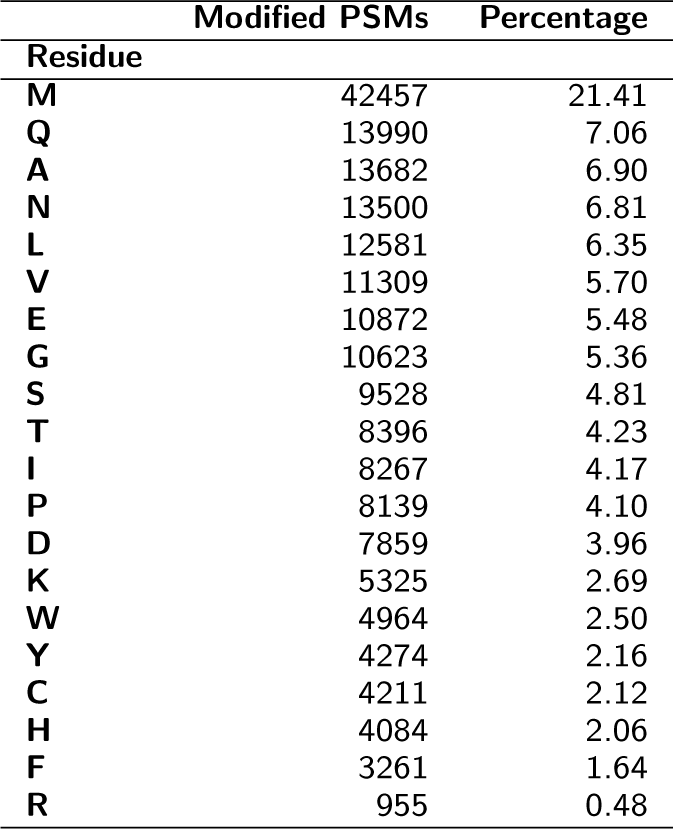
Most Commonly Modified AA Residues

| Residue | Modified PSMs | Percentage |
| --- | --- | --- |
| M | 42457 | 21.41 |
| Q | 13990 | 7.06 |
| A | 13682 | 6.90 |
| N | 13500 | 6.81 |
| L | 12581 | 6.35 |
| V | 11309 | 5.70 |
| E | 10872 | 5.48 |
| G | 10623 | 5.36 |
| S | 9528 | 4.81 |
| T | 8396 | 4.23 |
| I | 8267 | 4.17 |
| P | 8139 | 4.10 |
| D | 7859 | 3.96 |
| K | 5325 | 2.69 |
| W | 4964 | 2.50 |
| Y | 4274 | 2.16 |
| C | 4211 | 2.12 |
| H | 4084 | 2.06 |
| F | 3261 | 1.64 |
| R | 955 | 0.48 |

We constructed the distribution of targeted amino acids for each mass shift by counting occurrences of each mass shift-AA pair across all nine time points. We observed significant differences among mass shifts in preference for a single type (or, in some cases, groups) of amino acid residues; the +22 Da and +38 Da modifications, for example, show a broad distribution across AA types, while +16 Da and −2 Da show strong (though not exclusive) preference for methionine. To quantify these differences in AA distribution, we ranked mass shifts by the ratio: PSMs for most common AA / mean(PSMs for all other AAs). AA distributions for the the top ranked (most biased towards one AA across multiple biological replicates) mass shifts are shown in Figure 3.

**Figure 3.**
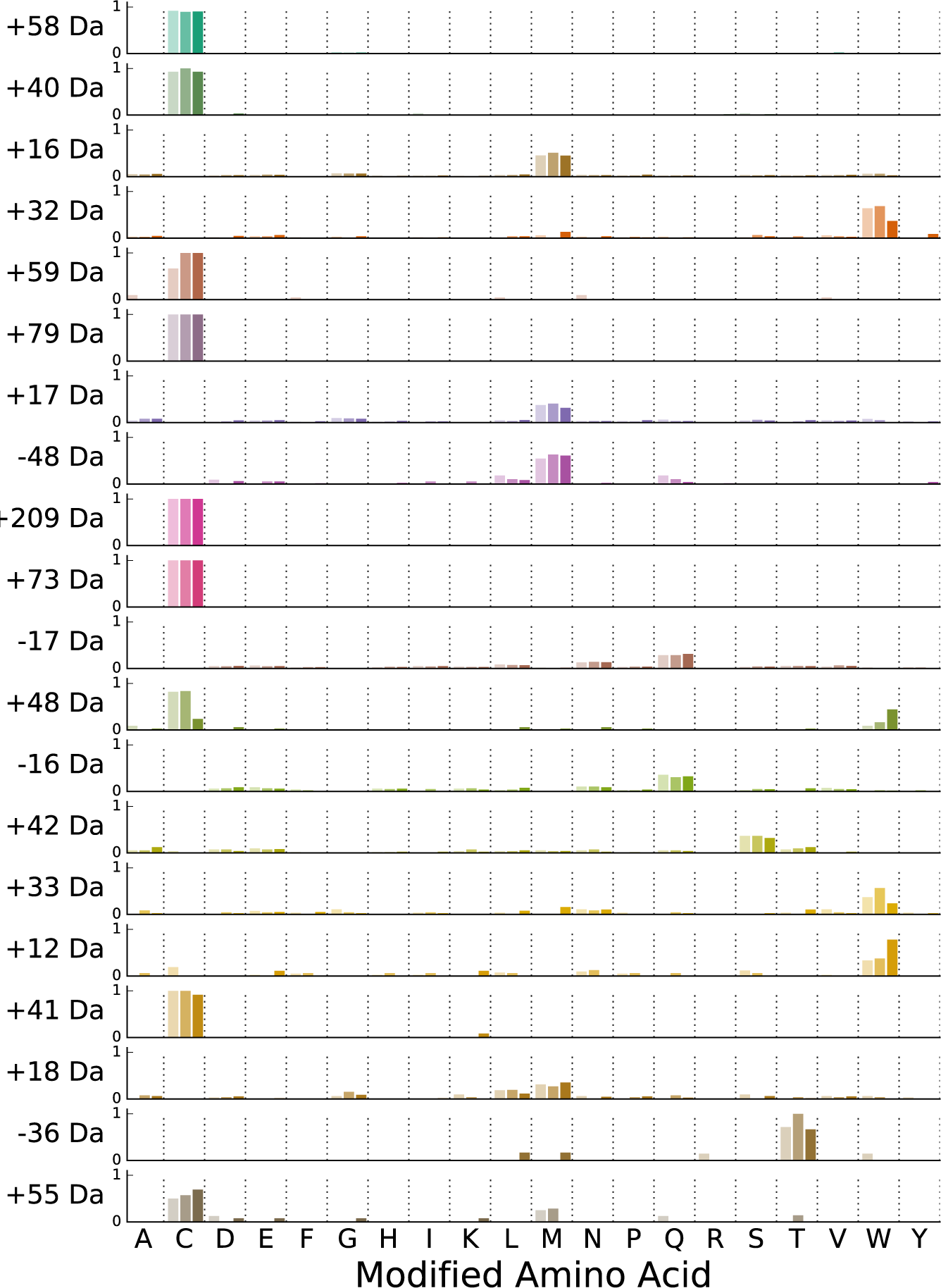
Distribution of selected mass shifts across amino acids. Height of bars within each row represents the fraction of total AA positions for the mass shift (indicated on the *y* axis) that were identified on each amino acid residue type (columns). Individual bars within each column represent fractions for each biological replicate (replicates 1,2, and 3 from left to right within each column). Mass shifts are ordered by the single-AA bias score (the ratio of counts for the most commonly modified AA type to the mean of the counts for all other types; see Methods) with the highest score (most biased for a single AA) at the top; only the top 20 mass shifts are shown. Note that a constant mass shift of +57 Da was added to all cysteine modifications to correct for the presence of carbamidomethylation, meaning that a small number of cysteine modifications (e.g. +209 Da) fall outside of the mass range scanned by MODa (+/− 200 Da).

A large number of modifications with a strong preference for cysteine residues were identified in all three biological replicates; most of these are likely artifacts of IAA treatment during sample preparation, and correspond to common modifications co-occuring with carbamidomethylation (+57 Da), e.g. +58 Da (57 + 1 Da), +59 Da (57 + 2 Da), and +40 Da (57 - 17 Da). The +209 Da mass shift, corresponding to a carbamidomethylated dithiothreitol modification of the cysteine thiol group, is a minor artifact of the reduction and alkylation of cysteine during sample preparation. The +48 Da mass shift was almost exclusively found at catalytic cysteine residues in a handful of proteins, and corresponds to the hyperoxidation of the cysteine thiol group (Cys-SH) into cysteic acid (Cys-SO_3_H). This modification is likely to be inactivating and irreversible, resulting in the increased accumulation of the modified form throughout the stationary phase. Among modifications targeting non-Cys residues, putative oxidative modifications show the strongest bias towards a single AA, with the +32 Da and +16/+17 Da modifications showing strong preferences for tryptophan and methionine, respectively. The eighth-ranked —48 Da modification is likely also a result of oxidation via dethiomethylation of methionine residues [42]. The strong preference of the acetylation mass shift (+42 Da) for serine is largely due to modifications on protein N-termini (see Section “N-terminal and C-terminal modifications”). A table of counts for each mass shift-amino acid pair is included in Additional File 12.

### N-terminal and C-terminal modifications

To search for modifications that preferentially occur at protein N and C termini, we used Fisher’s exact test (FET) to compare the ratio of modified : unmodified counts of each mass shift occurring at the N or C terminus of a protein to the same ratio for mass shifts occurring at all other positions. FET *p*-values for N-terminal and C-terminal enrichment were calculated for all mass shifts within each biological replicate and filtered for consistency by requiring all three replicates to have *p* < 0.05. Nt- and Ct-biased mass shifts are shown in Tables 4 and 5. We also examined the distribution of unique modified positions for these Nt- and Ct-biased mass shifts as a function of normalized protein length, to determine whether the observed positional bias was a general feature of the mass shift or due to a small number of highly abundant modified positions (Figures 4 and 5).

**Figure 4.**
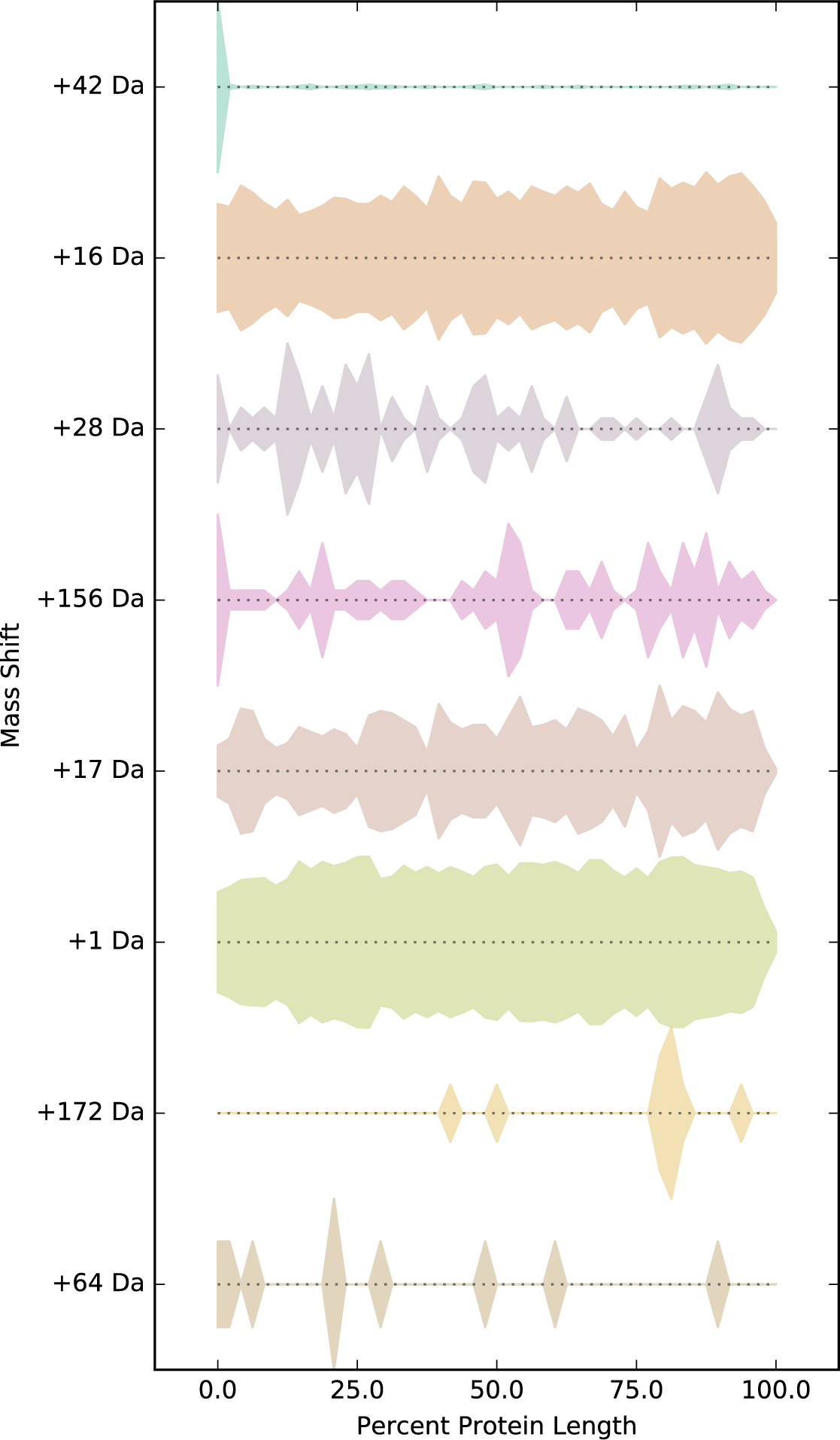
Distribution of Nt-biased mass shifts across positions in protein sequence. Width of traces within each row represent the density of unique positions identified for the mass shift indicated to the left along target proteins, normalized by protein length (*x*-axis). Traces are plotted symmetrically about the *x*-axis. Mass shifts are ranked from top to bottom by combined *p*-value from the Fisher’s exact test for N-terminal modification enrichment across all three replicates (see section “N-terminal and C-terminal Modifications” and Table 4), with mass shifts having the strongest N-terminal enrichment at the top.

**Figure 5.**
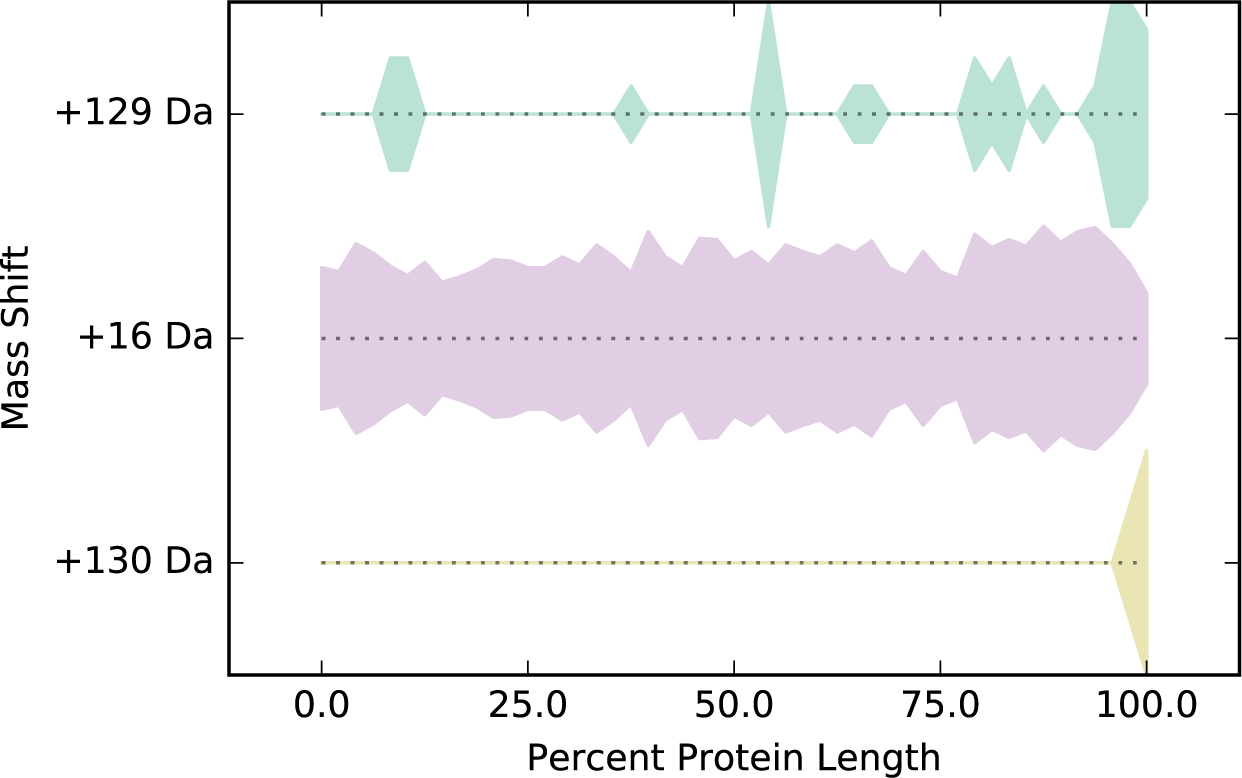
Distribution of Ct-biased mass shifts across positions in protein sequence. Width of traces within each row represent the density of unique modified positions (i.e. positions with more than one modified PSM; each position is counted once per protein) identified for each mass shift (indicated to the left) along target proteins, normalized by protein length (*x*-axis). Traces are plotted symmetrically about the *x*-axis. Mass shifts are ranked from top to bottom by combined *p*-value from the Fisher’s exact test for N-terminal modification enrichment across all three replicates (see section “N-terminal and C-terminal Modifications” and Table 5), with mass shifts having the strongest C-terminal enrichment at the top.

**Table 4.** Mass Shifts occuring more frequently on N-terminal ends of proteins

| Mass Shift | Biol. Replicate | Protein-N-terminal-Biased Mass Shifts |  | *p*-value |
| --- | --- | --- | --- | --- |
|  |  | N-terminal | Non-N-Terminal |  |
| **+42 Da** | 1 | 759 / 1869 (40.61%) | 132 / 7860 (1.68%) | 0.0 |
|  | 2 | 657 / 1561 (42.09%) | 111 / 6814 (1.63%) | 0.0 |
|  | 3 | 682 / 1930 (35.34%) | 147 / 7681 (1.91%) | 0.0 |
| **+16 Da** | 1 | 1187 / 8016 (14.81%) | 13633 / 510869 (2.67%) | 0.0 |
|  | 2 | 972 / 6971 (13.94%) | 9350 / 385682 (2.42%) | 0.0 |
| | 3 | 531 / 5216 (10.18%) | 20571 / 423856 (4.85%) | $7.36 \times 10^{-55}$ |
| **+28 Da** | 1 | 16 / 128 (12.50%) | 54 / 31261 (0.17%) | $4.56 \times 10^{-24}$ |
| | 2 | 27 / 136 (19.85%) | 11 / 28221 (0.04%) | $1.78 \times 10^{-55}$ |
| | 3 | 45 / 140 (32.14%) | 305 / 33420 (0.91%) | $1.65 \times 10^{-54}$ |
| **+156 Da** | 1 | 20 / 1051 (1.90%) | 277 / 54832 (0.51%) | $1.17 \times 10^{-6}$ |
| | 2 | 14 / 832 (1.68%) | 154 / 58376 (0.26%) | $1.34 \times 10^{-7}$ |
| | 3 | 26 / 674 (3.86%) | 313 / 43231 (0.72%) | $2.73 \times 10^{-11}$ |
| **+17 Da** | 1 | 60 / 3107 (1.93%) | 585 / 178417 (0.33%) | $8.94 \times 10^{-26}$ |
| | 2 | 45 / 2730 (1.65%) | 512 / 116280 (0.44%) | $6.38 \times 10^{-13}$ |
| | 3 | 54 / 2070 (2.61%) | 1960 / 147203 (1.33%) | $7.64 \times 10^{-6}$ |
| **+1 Da** | 1 | 138 / 6868 (2.01%) | 27945 / 2042671 (1.37%) | $1.88 \times 10^{-5}$ |
| | 2 | 151 / 5934 (2.54%) | 25111 / 1455121 (1.73%) | $5.28 \times 10^{-6}$ |
| | 3 | 204 / 6226 (3.28%) | 30808 / 1810086 (1.70%) | $1.52 \times 10^{-17}$ |
| **+172 Da** | 1 | 15 / 15 (100.00%) | 0 / 9431 (0.00%) | $3.11 \times 10^{-48}$ |
| | 2 | 1 / 1 (100.00%) | 1 / 11114 (0.01%) | $1.8 \times 10^{-4}$ |
| | 3 | 14 / 14 (100.00%) | 11 / 7365 (0.15%) | $2.77 \times 10^{-37}$ |
| **+64 Da** | 1 | 13 / 58 (22.41%) | 13 / 1620 (0.80%) | $1.84 \times 10^{-13}$ |
| | 2 | 4 / 48 (8.33%) | 6 / 1340 (0.45%) | $2.27 \times 10^{-4}$ |
| | 3 | 13 / 103 (12.62%) | 19 / 2119 (0.90%) | $3.58 \times 10^{-10}$ |

**Table 5.** Mass Shifts occuring more frequently on C-terminal ends of proteins

| Mass Shift | Biol. Replicate | C-terminal-Biased Mass Shifts |  | *p*-value |
| --- | --- | --- | --- | --- |
|  |  | C-terminal | Non-C-terminal |  |
| **+129 Da** | 1 | 165 / 703 (23.47%) | 78 / 10189 (0.77%) | $2.00 \times 10^{-142}$ |
| | 2 | 212 / 886 (23.93%) | 49 / 8541 (0.57%) | $1.98 \times 10^{-177}$ |
| | 3 | 82 / 502 (16.33%) | 59 / 8478 (0.70%) | $4.36 \times 10^{-67}$ |
| **+16 Da** | 1 | 39 / 205 (19.02%) | 14781 / 518680 (2.85%) | $7.88 \times 10^{-21}$ |
| | 2 | 42 / 245 (17.14%) | 10280 / 392408 (2.62%) | $7.76 \times 10^{-22}$ |
| | 3 | 22 / 103 (21.36%) | 21080 / 428969 (4.91%) | $5.13 \times 10^{-09}$ |
| **+130 Da** | 1 | 56 / 427 (13.11%) | 2 / 366 (0.55%) | $1.18 \times 10^{-14}$ |
| | 2 | 75 / 542 (13.84%) | 0 / 549 (0.00%) | $1.02 \times 10^{-24}$ |
| | 3 | 18 / 276 (6.52%) | 0 / 196 (0.00%) | $8.29 \times 10^{-05}$ |

Eight mass shifts were identified as Nt-biased after filtering (Table 4 and Figure 4). The strongest Nt preference is displayed by the +42 Da mass shift, corresponding to N-terminal acetylation, with modified N termini representing 35-42 % of total observed counts for positions with at least one +42 Da count. The remaining Nt-biased mass shifts fall into two broad categories. The first are rare modifications that occur at a small number of positions at high frequency, such as the +28 Da mass shift (possible retention of formylation on an Nt-terminal fMet, 12-32%), the +64 Da mass shift (possible modification by acetate, 8-22%) and the +172 Da mass shift (100% in all replicates). The second category is comprised of common modifications that occur at low frequency across a larger number of positions; this includes oxidation (+16 Da, 10-14%), most commonly of a retained Nt methionine, and hydrogenation (+1 Da, 2-3%). A beneficial feature of our analysis is the ability of MODa to identify modified N-terminal residues even in the presence of un-annotated N-terminal methionine cleavages. For the protein SecB, for example, we recovered abundant N-terminal peptides which had both undergone N-terminal Met cleavage and putative acetylation at the penultimate N-terminal Ser residue (see Additional File 12), despite the fact that this protein had not been annotated as having its N-terminal Met cleaved in the UniProt database.

Only three mass shifts were identified as Ct-biased after consistency filtering (Table 5 and Figure 5). Two of these, +129 Da (16-24% of counts at C-terminal positions across the three replicates modified, compared to < 1% of counts at all other positions) and +130 Da (6.5-14% of counts at C-terminal positions across the three replicates modified, compared to < 1% of counts at all other positions), most likely correspond to the same modification, C-terminal addition of a glutamate residue. Interestingly, the third C-terminal mass shift is oxidation (+16 Da), which is observed to occur at high frequency (17-20% modified counts across replicates at C-terminal residues with at least one +16 Da modification, compared to 2.6-5% at all other modified positions) on C-terminal residues as well as N-terminal residues, although the C-terminal modification is observed for a smaller set of proteins.

The C-terminal glutamylation modification is especially interesting. The most frequent target for this modification is the C terminus of the 30S ribosomal protein S6 (RpsF), which is known to undergo post-translational modification with 1-4 glutamate residues (mass = 129 Da) [43]. The enzymatic addition of these Glu residues to S6 proceeds in a stepwise fashion, and any modification of two or more Glu residues would fall outside the range of mass shifts that were considered in our analysis, so it is likely that the mono-glutamylated S6 we observed only represents a subset of the total modified S6 present in our samples.

We also identified a previously unreported C-terminal +129 Da modification of the stationary phase ribosomal stability factor RaiA / YfiA [22]. YfiA binds within the mRNA tunnel of the 30S subunit [44, 45], where it inhibits translation [44, 46] and prevents subunit dissociation and 100S dimer formation for a subset of ribosomes in stationary phase [47]. YfiA and S6 lie near one another within the 30S subunit, and both proteins’ C termini extend towards the same region of the 16S rRNA on the subunit surface (Figure S2), although the modified C-terminal tails themselves are not resolved in the crystal structure. The temporal modification patterns of S6 and YfiA differ dramatically (Figure S3). S6 levels of both total PSM counts and Ct +129-Da modified counts peak in mid-exponential phase, followed by a steep drop to a lower number of counts that is maintained through late stationary phase; the relative proportion of +129 Da modified counts remains nearly unchanged across all time points. In contrast, YfiA shows low or no counts of either modified or unmodified PSMs until the onset of stationary phase, when overall counts increase dramatically, accompanied by a low but constant level of C-terminal +129 Da modification through late stationary phase. The exponential phase enrichment we observed for the +129 Da mass shift is therefore due largely to changes in overall expression of its target proteins rather than differential modification.

### Temporal patterns

The glucose starvation dataset used in our analysis is unique in the wide range of timepoints (3h-336h) that were sampled. Changes in abundance during different phases of the growth cycle in liquid culture have been observed for individual PTMs, but an unbiased examination of temporal variation in the global PTM profile has not been performed in *E. coli.* To identify mass shifts with significant frequency changes over the growth cycle, we first pooled four of our nine time-point samples into exponential-phase samples (3h, 4h, 5h, and 6h, EXP) and four into stationary-phase samples (24h, 48h, 168h, 336h, STA). (We did not include the 8h sample in this analysis.) We then grouped counts across modified amino-acid positions by mass shift-AA pairs and compared the ratio of modified:unmodified counts at all modified positions in the EXP and STA pools using Fisher’s exact test (FET) [48]. Mass shift-AA pairs were called as significant if their FET p-values passed a false-discovery rate filter (< 5% FDR by the Benjamini-Hochberg step-down procedure) in all three biological replicates. Because we used a two-tailed test that was unable to determine the direction of enrichment (i.e., EXP > STA or EXP < STA), we subsequently divided significant mass shift-AA pairs into EXP > STA or EXP < STA groups using the FET log-odds score.

We identified only a single mass shift that consistently shows significantly higher levels of modification in exponential phase across all three biological replicates, a +16 Da modification of tryptophan (3.78–4.33% of total counts at modified positions across the three biological replicates have the mass shift in exponential phase, 1.60–1.69% in stationary phase, Table 6). The behavior of this mass shift differs slightly across the three biological replicates: in biological replicates 1 and 2, the +16 Da Trp modification shows a spike in abundance near the Exponential-Stationary phase transition (8h), followed by a drop to near zero by mid-stationary phase (48h), while replicate 3 shows a spike of enrichment earlier in exponential phase (4h) followed by a steep drop off at the 5h timepoint (Figure 6).

**Figure 6.**
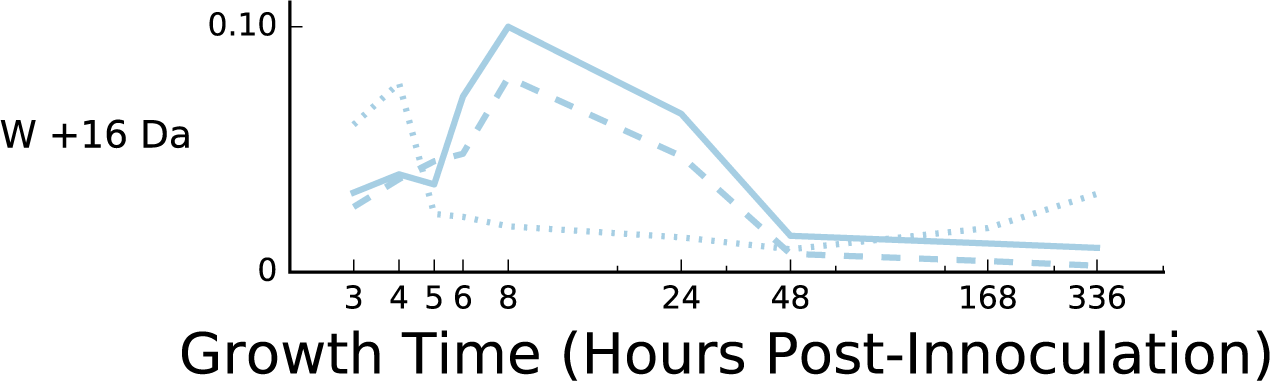
Abundance across all growth timepoints of tryptophan monooxidation, the sole mass shift with stronger modification in exponential phase. Plot shows the fractional modification *N*_mod_/(*N*_mod_ + *N*_unmod_) across all nine time points, for positions having at least one W +16 Da modification at any time point. Individual traces show results for individual biological replicates 1 (solid lines), 2 (dashed lines), and 3 (dotted lines).

**Table 6.** Mass-shift–amino-acid pairs with elevated frequency in exponential phase

| Mass Shift | Amino Acid | Biol. Replicate | Exponential | Stationary | *p*-value |
| --- | --- | --- | --- | --- | --- |
| +16 Da | W | 1 | 285/6836 (4.17%) | 248/8775 (2.83%) | $5.55 \times 10^{-06}$ |
| | | 2 | 238/6182 (3.85%) | 141/7242 (1.95%) | $4.31 \times 10^{-11}$ |
| | | 3 | 282/6182 (4.56%) | 113/6414 (1.76%) | $9.13 \times 10^{-20}$ |

We identified five mass shifts that consistently show significantly higher levels of modification in stationary phase across all three biological replicates: a +1 Da modification of asparagine (1.90-3.06% of total counts at modified positions have the mass shift in exponential phase, 2.95-4.60% in stationary phase); +42 Da modifications of serine, alanine, and threonine (29.78-31.71% EXP, 46.30-60.07% STA; 18.33-22.45% EXP, 34.81-46.46% STA; and 0.0-3.37% EXP, 9.46-15.70%, respectively), and a +48 Da modification of cysteine (0.94-1.09% EXP, 3.11-4.19% STA) (Table 7). As with the exponential-phase-biased mass shifts, we observed different temporal patterns when timepoints are considered individually (Figure 7). For example, the +1 Da asparagine modification and the +48 Da cysteine modification show steady increases across stationary phase, reaching their highest value at the latest stationary phase timepoint (336h), while the +42 Da modification to serine shows a more step-like increase in abundance near the onset of stationary phase, with abundance remaining fairly constant through the latest timepoints.

**Figure 7.**
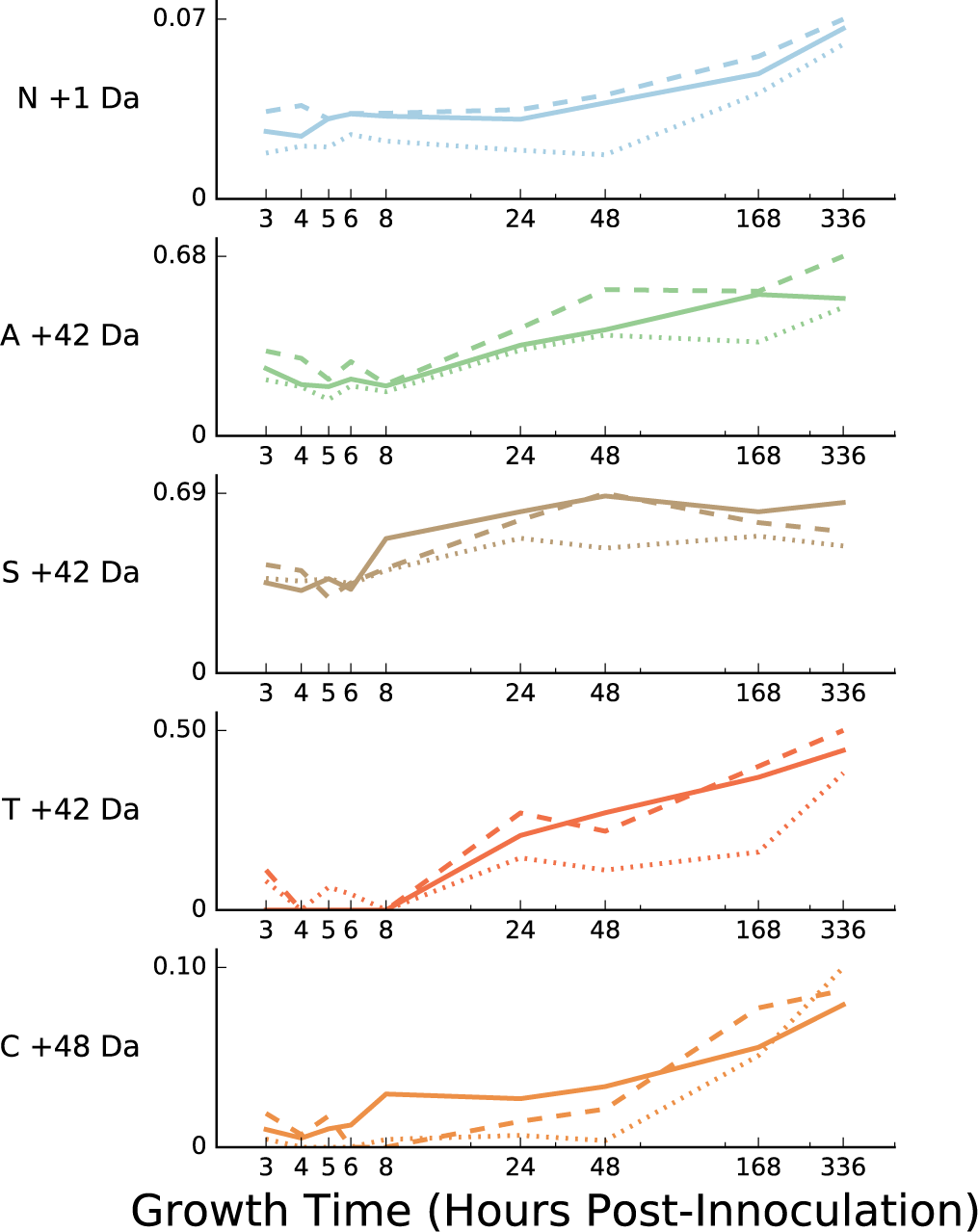
Abundance across all growth timepoints of mass shifts with stronger modification in stationary phase. Each plot shows the fractional modification *N*_mod_/(*N*_mod_ + *N*_unmod_) across all nine time points, for positions having at least one modification of the indicated type at any time point. Individual traces within each plot show results for individual biological replicates 1 (solid lines), 2 (dashed lines), and 3 (dotted lines). Mass shift are ranked from top to bottom by p-value from the Fisher’s exact test for modification enrichment in exponential phase (STA > EXP; see Section “Temporal patterns” and Table 7), averaged across all three replicates, with the most stationary-phase-enriched (lowest *p*-values) at the top.

**Table 7.**
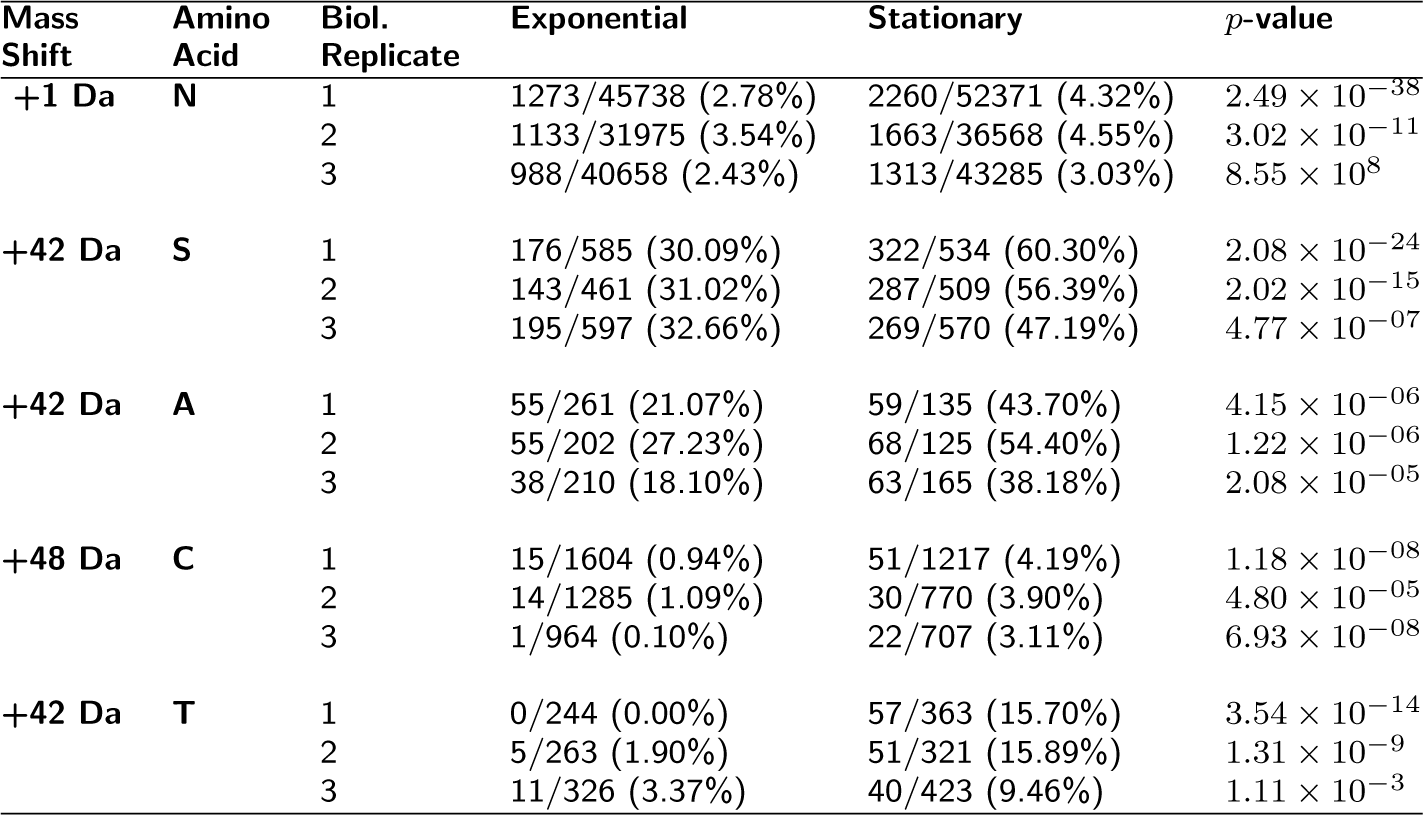
Mass-shift–amino-acid pairs with elevated frequency in stationary phase

| Mass Shift | Amino Acid | Biol. Replicate | Exponential | Stationary | *p*-value |
| --- | --- | --- | --- | --- | --- |
| +1 Da | N | 1 | 1273/45738 (2.78%) | 2260/52371 (4.32%) | $2.49 \times 10^{-38}$ |
| | | 2 | 1133/31975 (3.54%) | 1663/36568 (4.55%) | $3.02 \times 10^{-11}$ |
| | | 3 | 988/40658 (2.43%) | 1313/43285 (3.03%) | $8.55 \times 10^8$ |
| +42 Da | S | 1 | 176/585 (30.09%) | 322/534 (60.30%) | $2.08 \times 10^{-24}$ |
| | | 2 | 143/461 (31.02%) | 287/509 (56.39%) | $2.02 \times 10^{-15}$ |
| | | 3 | 195/597 (32.66%) | 269/570 (47.19%) | $4.77 \times 10^{-07}$ |
| +42 Da | A | 1 | 55/261 (21.07%) | 59/135 (43.70%) | $4.15 \times 10^{-06}$ |
| | | 2 | 55/202 (27.23%) | 68/125 (54.40%) | $1.22 \times 10^{-06}$ |
| | | 3 | 38/210 (18.10%) | 63/165 (38.18%) | $2.08 \times 10^{-05}$ |
| +48 Da | C | 1 | 15/1604 (0.94%) | 51/1217 (4.19%) | $1.18 \times 10^{-08}$ |
| | | 2 | 14/1285 (1.09%) | 30/770 (3.90%) | $4.80 \times 10^{-05}$ |
| | | 3 | 1/964 (0.10%) | 22/707 (3.11%) | $6.93 \times 10^{-08}$ |
| +42 Da | T | 1 | 0/244 (0.00%) | 57/363 (15.70%) | $3.54 \times 10^{-14}$ |
| | | 2 | 5/263 (1.90%) | 51/321 (15.89%) | $1.31 \times 10^{-9}$ |
| | | 3 | 11/326 (3.37%) | 40/423 (9.46%) | $1.11 \times 10^{-3}$ |

### Preferential persistence of N-terminally acetylated proteins in stationary phase

The N-terminal bias and preference for serine, alanine, and threonine residues observed for the +42 Da mass shift strongly suggests that this modification corresponds to N-terminal Nα-acetylation. Although cotranslational N-terminal Na acetylation (NtAc) is widespread in eukaryotic proteins, the prevalence and physiological significance of this modification in prokaryotes is poorly understood. In *E. coli,* only five native proteins are known to possess an NtAc modification: the ribosomal proteins S5 (encoded by the rpsE gene), S18 (encoded by the rpsR gene), and L12/7 (encoded by the rplL gene)[49]; elongation factor Tu (EFTu, encoded by the tufB gene) [50]; and the chaperone SecB [51]. In addition, a number of heterologous eukaryotic proteins are modified with an NtAc when overexpressed in *E. coli* [52, 53, 54, 55].

We identified more than 40 Nt-acetylated proteins, and were able to recover modified peptides from known Nt-acetylation target SecB (Figure S4) and a small number of peptides matching Nt-acetylated ribosomal protein S5 (Figure S6) in our initial MODa dataset. The low peptide counts for S5, as well as the absence of modified PSMs for the other known (and highly abundant) targets ribosomal proteins S18 and L7/12, as well as EFTu, are likely due to the presence of tryptic cleavage sites within a few residues of the N-terminus in all three of these proteins (Nt-AHIE**K** QAGE for S5, Nt-A**R**YF**RRRK**F for S18, Nt-SIT**K**DQIEE for L7/12, and Nt-S**K**E**K**FERT**K** for EFTu). This means that most copies of the protein present in our samples will produce N-terminal peptides too short to recover during subsequent liquid chromatography and MS/MS steps. Consistent with this interpretation, we were able to recover abundant peptides from non-N-terminal regions of all four of these proteins, and the small number of S5 N-terminal peptides that were recovered were all the result of missed cleavage events at the N-terminal-most cleavage site. Among the NtAC peptides that were recovered in our modA dataset, the Nt fragment from SecB is by far the most frequently observed, representing 15–41% of the total Nt-Acetylated peptides across the nine time points. In addition, six other proteins from our dataset were previously identified as Nt-acetylation targets in an enrichment-based analysis of N-terminal modifications in *Pseudomonas aeruginosa* [13] (See Table 8).

**Table 8.**
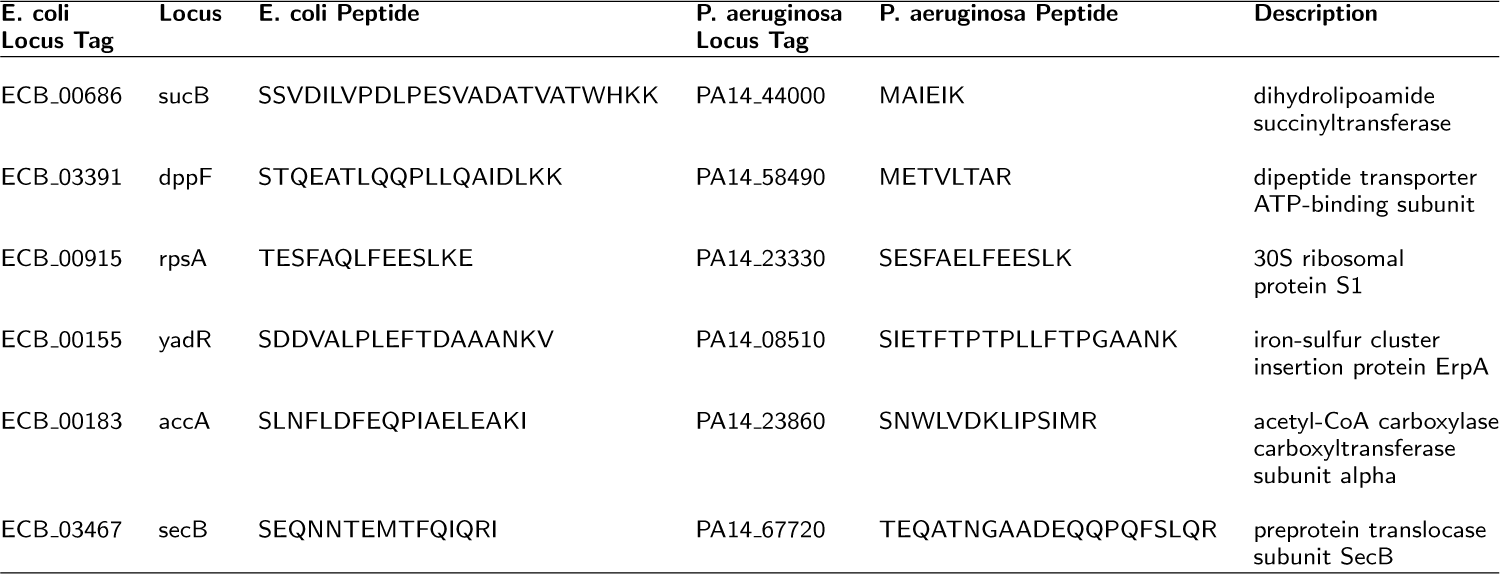
Overlapping N-terminal Nα-acetylation targets between current data and *P. aeruginosa* [13]

We observed that NtAc modified proteins are proportionally more heavily modified in stationary phase (Figure 7). This pattern could be explained by (i) an increase in acetylation activity in stationary phase and/or (ii) a proportionally larger decrease in non-acetylated copies of a protein relative to acetylated copies in stationary phase. To differentiate between these scenarios, we plotted total PSM counts and NtAc-modified PSM counts for pooled NtAc-targeted proteins across all nine time points (Figure 8). When all NtAc-targeted proteins are considered (Figure 8, top left panel), the total number of PSMs stays appoximately constant, while the number of NtAc-modified PSMs increases by nearly twofold in early stationary phase, consistent with scenario (i). However, NtAc-targeted proteins pooled by penultimate amino acid (Figure 8) or individual NtAc-targeted proteins (Figures S4, S5, and S6) show a mixture of both scenarios. NtAc-targeted proteins with a penultimate serine or threonine residue, for example, exhibit a pattern consistent with scenario (i), similar to the pattern for all targets (Figure 8, top right and bottom left panels). Proteins with a penultimate alanine, however, show a slight increase in modified peptides at the onset of stationary phase, accompanied by a large drop in unmodified peptides (Figure 8, top right panel). Many of the the most heavily NtAc-modified proteins also show this pattern, such as LysS, SpeA, pdxH, and SecB (Figure S4), and IlvA and KdgR (Figure S4). This preferential retention of NtAc-modified peptides in stationary phase suggests that NtAc may play role in protein stability by acting as an anti-degradation signal (see Discussion). A table of all Nt-acetylation sites recovered by MODa is included in Additional File 12.

**Figure 8.**
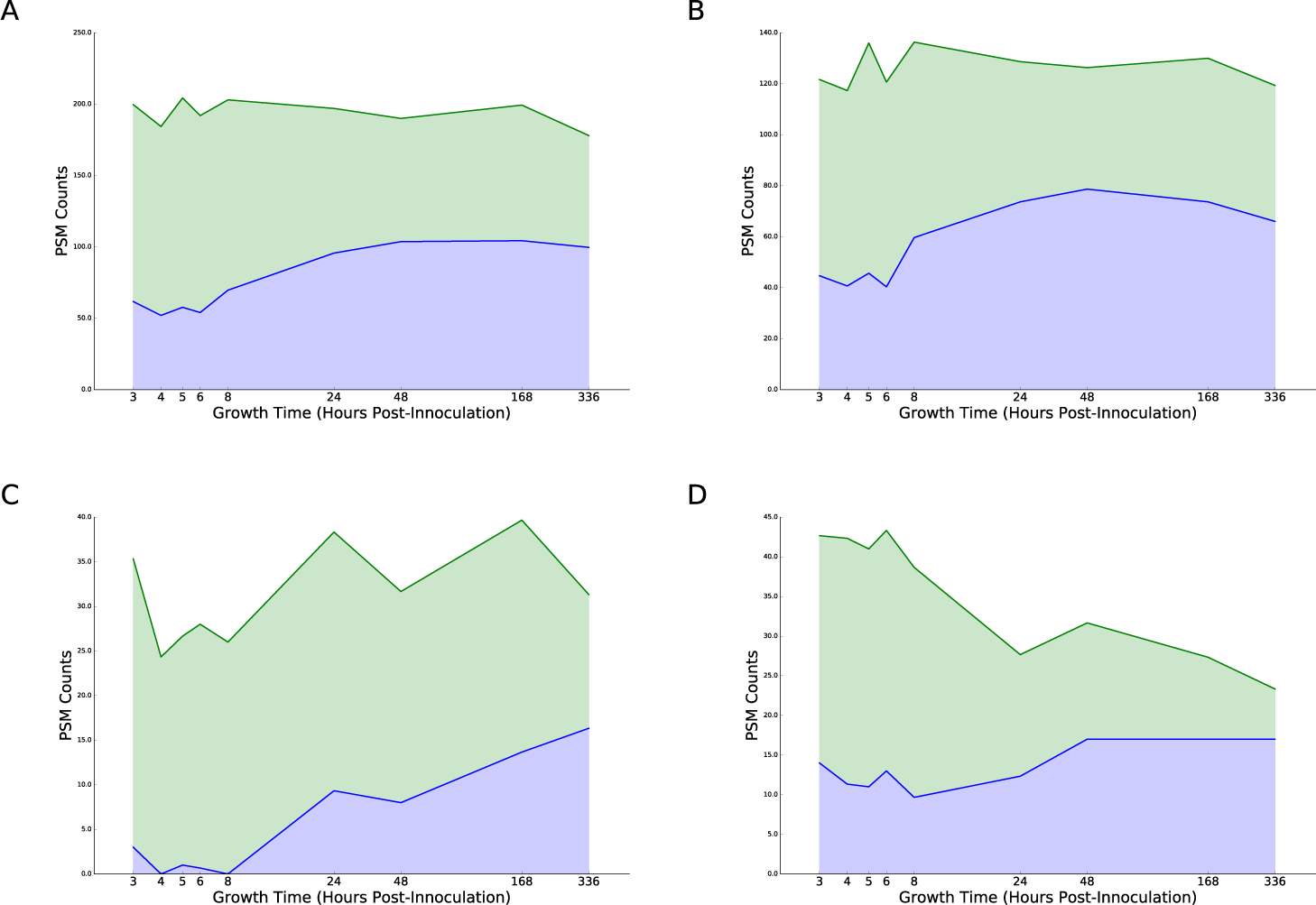
N-terminal +42 Da modified proteins are preferentially retained in stationary phase. Plots show unmodified (green) and +42 Da modified (blue) PSM counts for all N-terminal positions possessing at least one +42 Da modification at any time point, averaged across the three biological replicates. Shown are total counts (A), counts for peptides with a penultimate (i.e. following a cleaved N-terminal methionine) serine residue (B), a penultimate threonine residue (C), and a penultimate alanine residue (D).

### Asparagine deamidation is strongly enriched in very late stationary phase

An interesting temporal pattern was also identified for the +1 Da modification of Asparagine residues, which increases in frequency throughout stationary phase and peaks at the last timepoint (336h) (Figures 7 and S7). +1 Da modifications were the most frequently observed modifications in our dataset, and likely result from a variety of different modification chemistries. A +1 Da modification occurring on an asparagine residue, however, is known to be a signature of nonenzymatic asparagine deamidation, in which a backbone nitrogen initiates a nucleophilic attack on the amide carbon of the asparagine side chain (or the asparagine amide nitrogen on the backbone carbonyl carbon) to form a cyclic succinimide intermediate [56, 57, 58]. This intermediate can then resolve by hydrolysis to either convert the original asparagine to an aspartate residue, or rearrange to form an isopeptide linkage through isoaspartate; both of these events result in a +1 Da mass shift.

Although asparagine deamidation can occur spontaneously as an experimental artifact during preparation of proteomic samples [59], a number of lines of reasoning suggest that at least a subset of the modifications we observe were present in the samples prior to processing. First, we observe a nearly identical pattern of increasing Asp +1 Da modification across all three of our biological replicates (Figure 7). All timepoints were collected from a single set of cultures started on the same day, each biological replicate was grown independently (on a different day) from the others, and all timepoints from a single replicate were processed for proteomic analysis in parallel. The bulk of nonenzymatic deamidation during proteomic sample prep has been shown to occur during tryptic digest [59], with both longer incubation time and basic pH increasing the occurrence of deamidated peptides. Although no special steps were taken during preparation of the samples included in our analysis, both of these variables should be identical across all timepoints (because samples were processed together), and any nonenzymatic deamidation should therefore also be constant across timepoints. The pattern does not appear to be explained simply by increased expression of modified proteins during stationary phase, as the pattern is observed even for individual modifications that have high abundance in both exponential and stationary phases (Figure S7). A table of all putative asparagine deamidation sites recovered by MODa is included in Additional File 12.

### Oxidative modifications of methionine and tryptophan are variable across biological replicates

Oxidation (+16 Da) modifications, particularly of methionine, are very common in our data, but with the exception of +16 Da modification of tryptophan residues (Figure 6), oxidations in general are not identified as having a significant bias for either exponential or stationary phase. Both Met +16 Da and Trp +16 Da show significant variability among the three biological replicates, with replicates one and two showing a similar pattern of relative modification enrichment over time, while replicate three has a different pattern (Figures S10 and S8). In addition, for both modifications replicates one and two show a peak of modified peptide counts centered at or near the 8 h time point (the exponential-stationary phase transition; this timepoint was excluded from our initial comparisons of stationary vs. exponential enrichment), with the proportion of modified PSMs then decreasing to early-exponential-phase levels or below by 24 h.

The reason for the discrepancy between the third replicate and the two others is unclear; this observation in combination with the common occurrence of oxidative modifications as experimental artifacts [60] makes it difficult to draw any biological conclusions from the temporal patterns of oxidative modifications. We observe the discrepancy among replicates only for oxidative modifications and not for other modified peptide counts or overall peptide levels, so one possibility is that a difference in redox conditions in sample processing influenced the number of oxidized peptides that were recovered. Differential modification in the third replicate is apparent in the temporal modification patterns of individual target sites (Figures S9 and S11), but does not display a consistent pattern across sites. Tables of all methionine and tryptophan oxidation sites are included in Additional File 12.

## Discussion

We have leveraged a large proteomics dataset [39] and the fast multi-blind spectral alignment algorithm MODa [38] to construct a comprehensive, unbiased map of all protein post-translational modifications between −200 and +200 Da at 9 timepoints, spanning early exponential phase (3h post-innoculation) through late stationary / starvation phase (336h, or 2 weeks post-innoculation). From this map, we have identified post-translational mass shifts with statistically significant differences in modification stoichiometry between N- and C-terminal ends of proteins and between exponential and stationary phases. This analysis has enabled us to identify previously unobserved temporal patterns and novel target proteins for known modifications, and to identify possible novel modifications. Finally, by comparing temporal patterns of modified and unmodified PSM counts for individual AA positions, we have been able to identify a possible relationship between post-translational modification and protein degradation rate in stationary phase.

Although decades of work have been dedicated to studying the biochemical and physiological function of post-translational modifications, much of this work has focused on a handful of modification chemistries such as Ser/Thr phosphorylation and Lys acetylation. Technical limitations in instrument sensitivity, sample preparation, and data analysis have meant that even these well-studied PTMs are often studied in isolation, and their place in the overall context of the cell, in terms of the overall set of pathways and proteins that utilize them, their interaction with other modifications, and their abundance relative to other modifications, is lost. An intriguing feature of our dataset is the relative scarcity of the most commonly studied regulatory modifications, such as phosphorylation and acetylation; in the few cases where such modifications are identified, they tend to occur at low frequency, even on very abundant proteins (Table 2).

By contrast, we have identified examples of abundant modification for N-terminal acetylation and C-terminal glutamylation. While further work is necessary to establish that the orders-of-magnitude differences in abundance between these modifications and more well-known regulatory modifications reflects their actual abundance in the cell, our findings do suggest that these modifications may play a more important physiological role than previously thought. Both of these modifications are known to be installed in a regulated and specific pattern on ribosomal proteins, but their function either in the ribosomal context or on other targets is largely unknown. In eukaryotic cells, N-terminal (Nt) acetylation has a variety of functions, including regulating protein stability, ER trafficking, protein complex formation, and membrane attachment [61], but there is no evidence for a similar role in prokaryotic cells. Nt acetylation of *E. coli* 30S ribosomal subunits S5 and S18 is thought to affect 30S ribosomal assembly by governing direct contacts with the rRNA [62], but no function for prokaryotic Nt acetylation outside of the ribosome has been proposed. While Nt acetylation of eukaryotic proteins can either inhibit [63] or enhance [64] degradation rates, our evidence suggests that Nt-acetylated proteins in *E. coli* are subject to lower levels of degradation than their unmodified counterparts. The viability of mutants in the three known *E. coli* Nt-acetyltransferase enzymes, RimI, RimJ, and RimL [49], should make experimental investigation of this hypothesis a tractable and interesting avenue for future research.

Similarly, the physiological role of C-terminal (Ct) glutamylation has only recently begun to be uncovered. Early investigations identified a Ct glutamyltransferase enzyme, RimK, that installs poly-E tails on ribosomal protein S6 *in vivo* and *in vitro*[65], but the only phenotypic effect observed in *E. coli rimK* mutant strains (other than loss of S6 glutamylation) is increased resistance to the aminoglycoside antibiotics streptomycin, neomycin, and kanamycin [66, 67]. Nonetheless, RimK, and presumably S6 Ct glutamylation, are conserved across a wide range of bacterial species [68], and recent work in *Pseudomonas* found profound changes in proteome composition and compromised colonization and virulence phenotypes in Δ*rimK* strains [69]. Our novel finding of an additional target of C-terminal glutamylation, the ribosomal hibernation factor YfiA, offers an additional experimental handle with which to examine the biological and molecular functions of this modification. The association of both Ct-glutamylation target proteins with the ribosome is especially interesting, because some evidence suggests that RimK modifies S6 C-termini specifically on intact ribosomes [68, 70], and RimK is known to catalyze poly-L-glutamine formation in the absence of S6 [71]. The C-terminal amino-acid residues of YfiA resemble those of S6 only in the presence of two glutamate residues in the last two positions (DDAEAGDSEE for S6 and ANFVEEVEEE for YfiA), indicating that targeting may largely be a function of YfiA’s structural association with the ribosome rather than due to a specific sequence signal.

The presence of a gradual increase in asparagine deamidation throughout our stationary phase samples is an intriguing observation. Asparagine deamidation/isomerization events occur spontaneously at a low frequency at specific protein residues with favorable local structure and sequence context [57, 72], and they are often observed in proteins that undergo a low frequency of turnover such as muscle fiber proteins [73] and lens crystallins [74]. This clock-like behavior of Asp deamidation is consistent with our observation of a steady accumulation of the Asp +1 Da mark through very late stationary phase (336h), and it suggests that proteins having this modification have been retained with little or no turnover throughout stationary phase. Remarkably, many of the most heavily modified target proteins are part of large supra-molecular complexes, including six on ribosomal proteins (N113 and N544 of ribosomal protein S1, encoded by the *rpsA* gene, N77 and N146 of ribosomal protein S5, encoded by the *rpsE* gene, and N89 of ribosomal protein L14, encoded by the *rplN* gene), N64 of EFTu, N77 of the genomic DNA structural protein H-NS [75], and two positions (N110 and N111) on SucB, the E2 subunit of the 2-oxoglutarate dehydrogenase multienzyme complex (OGDHC) [76]. Although retention of intact ribosomes through stationary phase is a well-documented phenomenon [22, 23], and H-NS has been shown to be involved in late-stationary-phase survival [77], retention of the OGDHC complexes has not been previously observed.

Our work has several limitations. First, although the consistent temporal signal across multiple replicates strongly indicates that the major modifications discussed above are of biological origin, we cannot rule out the possibility that a subset of these modifications are experimental artifacts; the oxidative modifications and asparagine deamidation in particular are known to occur as artifacts of downstream sample processing in MS/MS [78, 60], so further experimental verification will be needed to confirm their biological origin. Our study is also limited by the need to examine a relatively small window of mass shifts (−200 Da to +200 Da); many known modifications fall outside of this window, such as glycosylation, longer chain acylations, and lipidations [79]. Finally, while the lack of bias for particular modifications offers a number of advantages in our analysis strategy, it also means that our results are more limited by the inherent sensitivity of both shotgun MS/MS and computational identification of PTMs. Consequently, our data are biased towards highly abundant proteins and mass shifts, a factor that likely explains the scarcity of well-known PTMs such as lysine acetylation and phosphorylation in our data. Equipment improvements and/or novel experimental procedures (e.g. [80]) will likely be needed to enable detection of low-abundance or short-lived PTM and other rare effects such as translational mutations [81].

## Conclusions

In summary, the work presented here highlights the holistic perspective and novel biological insights that can be generated by combining unbiased PTM detection and deep temporal sampling of bacterial growth. Stationary phase biology and post-translational modification in prokaryotic systems are both still areas of active research with many open questions, and we hope that the analysis paradigm presented here can be applied to additional organisms and growth conditions to gain broader insight into prokaryotic physiology and evolution.

## Methods

### Origin of the analyzed data

All data were taken from a previously published *E. coli* time course [39]. In that study, *E. coli* was grown in glucose minimal media and samples were collected at 8 different time points: 3, 4, 5, 8, 24, 48, 168 and 336 hours past inoculation. The entire experiment was carried out in triplicate, with cultures in each time course grown at different times. Mass-spectrometry on these samples was carried out as follows [39]: Protein samples were prepared by trypsin digest and each sample was then analyzed using liquid chromatography mass spectrometry (LC/MS) on a LTQ-Orbitrap (Thermo Fisher). The resulting data are available from the ProteomeXchange Consortium (accession PXD002140) [82].

### Post-translational modification identification and analysis

We analyzed the raw mass-spectrometry data via MODa [38]. MODa is a naive based spectral alignment algorithm that identifies peptides and their associated PTMs from the input mzXML spectral files. The program needs a few additional parameters, such as enzyme used, instrument used to capture the mass-spec data, precursor and product ion mass tolerances, fixed modifications, any rules to apply on the digest, such as semi-tryptic or fully-tryptic, number of modifications per peptide, and the mass-range to search for PTMs. We ran separate MODa searches for each of the 9 time points. Since there were 3 biological replicates, this resulted in a total of 27 MODa searches. We set the enzyme used in the searches to trypsin, with fully-tryptic and no-proline rules. We allowed for 2 missed cleavages. We used a mass-tolerance for the precursor ion of 10 ppm, and the mass-tolerance used for the product ion was set to 0.5 Da. Finally, we set carbamidomethylation (+57 Da) of cysteine as a static or fixed modification. As mentioned earlier, MODa requires a mass range to search for variable modifications, so we ran MODa searches for the mass range between −200 and +200 Da. We used the *E. coli* B REL606 genome sequence (GenBank:NC_012967.1 [83]) to create the reference proteome. For the independent analysis of N-terminal acetylation, a modified *E. coli* B REL606 database was constructed in which N-terminal methionines were removed for 359 proteins annotated in UniProt for cleavage of “initiator methionine”. Raw spectral data were searched against this database by SEQUEST-HT (Proteome Discoverer 1.4; Thermo Scientific). Only fully-tryptic peptides were considered, allowing up to two missed cleavages. A precursor mass tolerance of 10 ppm and fragment mass tolerance of 0.5 Da were used. Oxidized methionine and N-terminal acetylation were considered as dynamic modifications; carbamidomethylation of cysteine was selected as a static modification. High confidence PSMs were filtered at a false discovery rate of ! 1% as calculated by Percolator (q-value j 0.01, Proteome Discoverer 1.4; Thermo Scientific).

### FDR calculations using MODa probabilities

For each PSM assigned to a spectra by MODa, the algorithm calculates a probability P_MODa_ using a logistic regression model that uses a variety of spectral features as parameters, trained on a standard set of correct and incorrect spectral matches [38]. To restrict our dataset to only high-quality PSMs, we used this probability to estimate the False-Discovery Rate (FDR) of incorrect matches in our dataset by (i) ranking all PSMs by their P_MODa_ values, and then (ii) iteratively adding PSMs, starting from the highest-probability matches, and calculating the FDR as

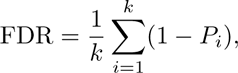

where *k* is the rank index of the last added PSM and *P*_*i*_ is the P_MODa_ of the ith ranked PSM, until adding any additional PSMs would result in an FDR above the chosen cutoff value.

### Metrics and statistical tests for single amino-acid bias, N-terminal/C-terminal bias, and growth-phase bias

To test the preference of each mass shift for modification of a single type of amino acid, we calculated a single-AA bias score *B_s_*(*A*) for mass shift s and length 20 vector A of counts of unique positions bearing at least one modification matching s for each amino acid type:

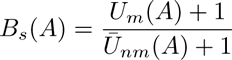

where *U_m_*(*A*) = max(*U*_*a*∈*A*_) and 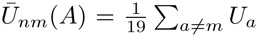. Note that “unique position” means that a given position in a protein is counted at most once regardless of total PSM counts at that position; this choice was intended to reduce bias from modifications with high abundance at a small number of positions.

To simplify our analysis, we constructed an intermediate dataset of PSM counts calculated by amino-acid position across all proteins in the REL606 annotated proteome. Unmodified counts *n*_*p*,unmod_ for each position *p* (having at least one modified or unmodified PSM) were calculated by summing PSM counts for any peptides that overlap *p* but do not have a modification (of any mass shift) at *p*. Modified counts *n_p,s_* were calculated by summing PSM counts for any peptides with a modification of mass shift *s* at protein position *p*.

To test for higher fractional modification by specific mass shifts at the protein termini, we constructed 2 × 2 contingency tables of the form

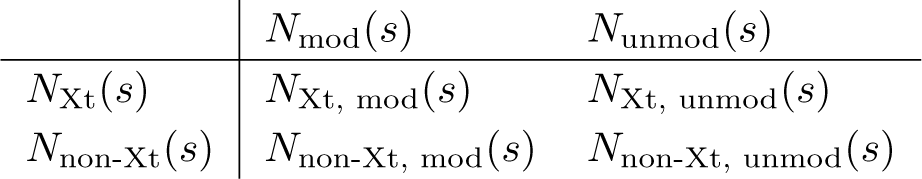

for each mass shift *s* in each of the three biological replicates, where *N*_xt_(*s*) is the sum Σ_*p*=Xt_(*n*_*p,s*_ + *n*_*p*,unmod_) for positions having at least one PSM with mass shift *s* occuring at the terminus Xt (either C- or N-terminus of a protein); *N*_non-Xt_(*s*) is the sum Σ_*p*≠Xt_(*n*_*p,s*_ + *n*_*p*,unmod_) for positions *p* having at least one PSM with mass shift *s*, occuring at all other positions (including the opposite terminus); *N*_mod_(*s*) is the sum Σ_*p,s*_ for positions having at least one PSM with mass shift *s*; and *N*_unmod_(*s*) is the sum Σ*n*_*p*,unmod_ for positions having at least one PSM with mass shift s. We used these tables to perform Fisher’s exact tests using a two-sided alternative hypothesis, implemented in Python using the statistics module of NumPy [84].

Similarly, to test for higher fractional modification by specific mass shift × amino acid pairs in either exponential or stationary phases of growth, we constructed 2 × 2 contingency tables of the form

| | $N_{\text{mod}}(s, a)$ | $N_{\text{unmod}}(s, a)$ |
| --- | --- | --- |
| $N_{\text{STA}}(s, a)$ | $N_{\text{Sta, mod}}(s, a)$ | $N_{\text{STA, unmod}}(s, a)$ |
| $N_{\text{EXP}}(s, a)$ | $N_{\text{Exp, mod}}(s, a)$ | $N_{\text{EXP, unmod}}(s, a)$ |

for each mass-shift-amino-acid pair in the three biological replicates, where *N*_mod_(*s, a*) and *N*_unmod_(*s, a*) are as above and *N*_EXP_(*s, a*) is the sum Σ_*t*=3,4,5,6_(*n*_*p,s,t*_+*n*_*p*,unmod,*t*_), where *n*_*p,s,t*_ is the count of PSMs modified by mass shift *s* at position *p* in timepoint *t* for positions *p* of amino acid type *a* having at least one PSM with mass shift *s*; and *N*_STA_(*s,a*) is the sum Σ_*t*=24,48,168,336_(*n*_p,s,t_ + *n*_*p*,unmod_,*t*) for positions *p* of amino acid type *a* having at least one PSM with mass shift *s*. We used these tables to perform Fisher’s exact tests using a two-sided alternative hypothesis, implemented in Python using the statistics module of NumPy [84].

### Raw data and analysis scripts

All raw data and analysis scripts are available online in the form of a git repository at https://github.com/wilkelab/EcolWPTMs. The analysis was performed in iPython [85] notebooks using the NumPy and SciPy libraries [84] for numerical calculations, the Pandas library [86] for data processing, and the MatPlotLib library [87] for plotting. Macromolecular structures in Figure S2 were assembled in MacPyMOL (version v1.7.4.4; Schrödinger, LLC).

## Author’s contributions

Conceived and designed the experiments: C.W.B., V.S., D.R.B., E.M.M., J.E.B., and C.O.W. Performed the experiments: C.W.B. and V.S. Analyzed the data: C.W.B., V.S. Wrote the paper: C.W.B., V.S., D.R.B., M.D.P., E.M.M., J.E.B., and C.O.W.

## Acknowledgements

This project was funded by Army Research Office Grant W911NF-12-1-0390, National Institutes of Health Grant R01 GM088344, Welch Foundation Grant F-1780, and CPRIT Grant RP110782. EMM acknowledges additional funding from the National Institutes of Health (DP1 OD009572), National Science Foundation (IOS1237975), and the Welch Foundation (F-1515).

We thank John Houser and Kevin Drew for helpful discussions. The Texas Advanced Computing Center (TACC) provided high-performance computing support.

## Additional Files

**Figure S1.**
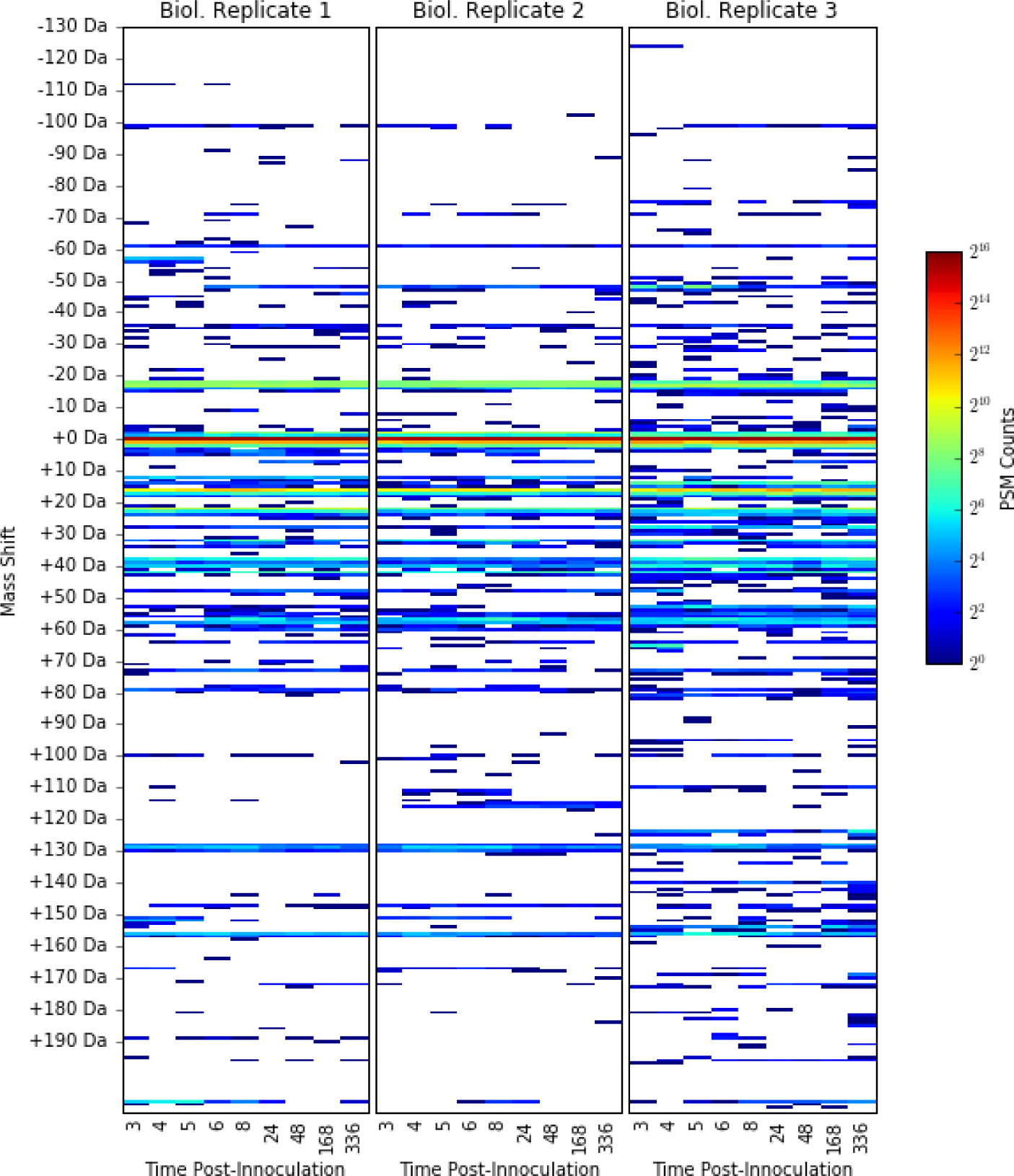
Abundance of all observed mass shifts across all 9 timepoints and 3 biological replicates. Color of heatmap corresponds to the log_2_-transformed count of MOD_a_-called modified PSMs in the 1% FDR set bearing the mass shift indicated on the *y*-axis for each of the nine timepoints (*x*-axis), for biological replicates 1, 2, and 3 (left, center, and right panels respectively). Although the MOD_a_ analysis was conducted for the mass window from −200 to +200 Da, no modifications were identified with mass shifts below −130 Da or above +196 Da.

**Figure S2.**
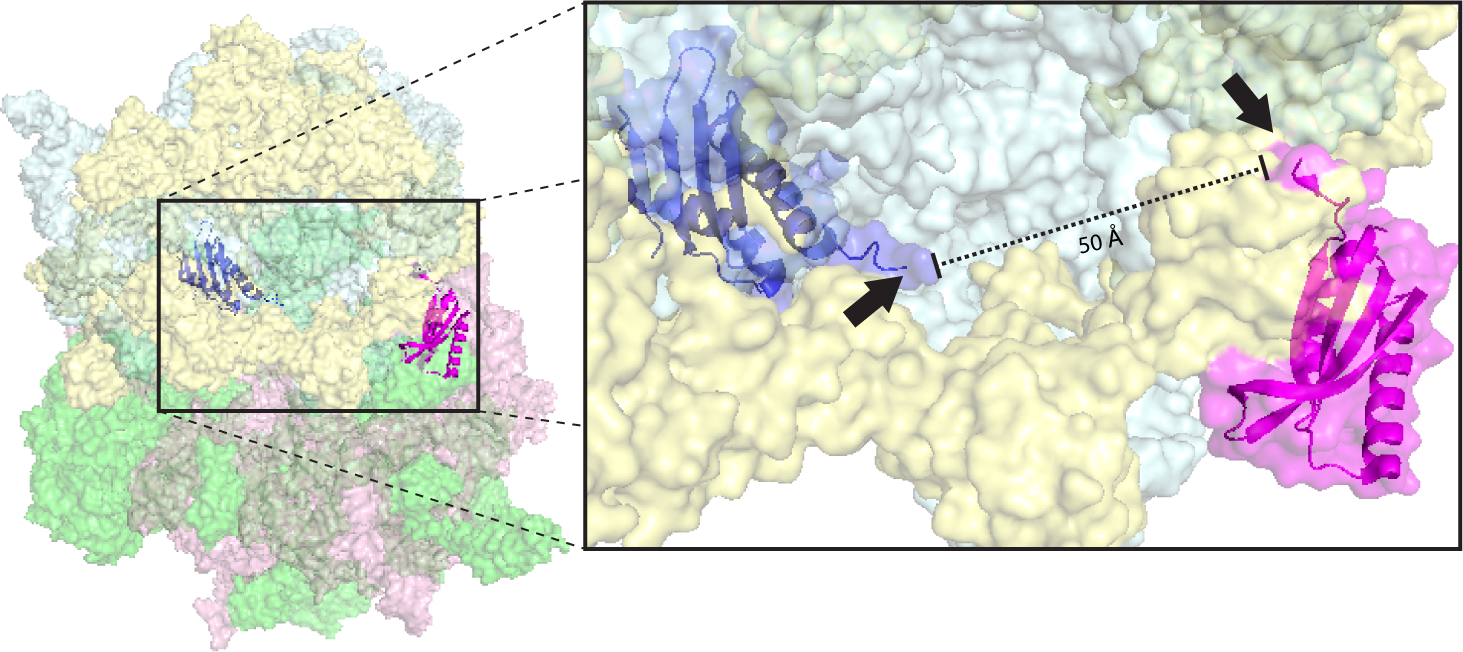
Location of YfiA and S6 in the 70S ribosome. Relative locations of YfiA (blue) and native *T. thermophilus* S6 (magenta) proteins in crystal structure of *E.coli* YfiA bound to the *T. thermophilus* 70S ribosome (PDB ID 4V8I [45]). YfiA is positioned within the 30S subunit mRNA tunnel, and S6 on the outer surface of the 30S subunit; the C-terminal tails of both proteins (black arrows) point toward the same region of the 16S rRNA (light blue). The 17 C-terminal residues for YfiA, including the terminal glutamate residues, were not resolved in the crystal structure; the *T. thermophilus* S6 protein coding sequence ends at residue 101, lacking the 30-AA unstructured C-terminal domain present in *E.coli* S6. 16S rRNA is shown in light blue; 30S ribosomal proteins (other than S6) are shown in light yellow; 50S ribosomal proteins are shown in green; and 23S rRNA is shown in pink.

**Figure S3.**
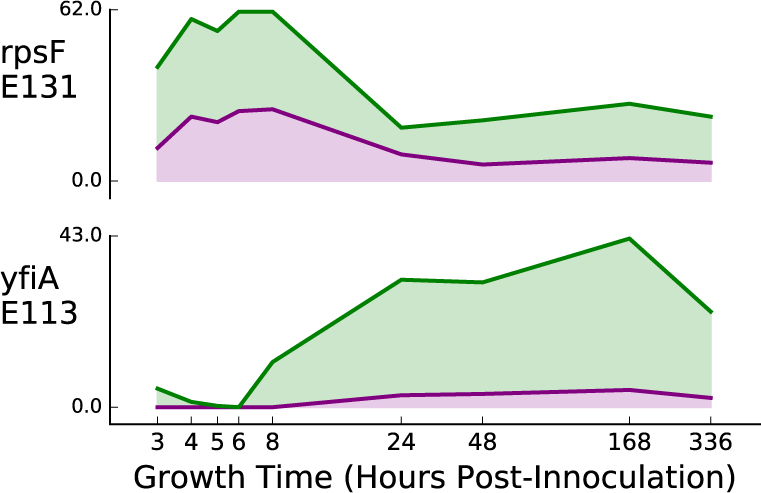
Modified and unmodified PSM counts for each AA position with a C-terminal +129 Da modification across all timepoints. Plots show unmodified (green) and +129 Da modified (purple) PSM counts across all nine timepoints (*x*-axis) for the C-terminal position of the two proteins that have at least one PSM identified by MODa as containing a C-terminal +129 Da modification. Counts represent the average of the three biologcial replicates.

**Figure S4.**
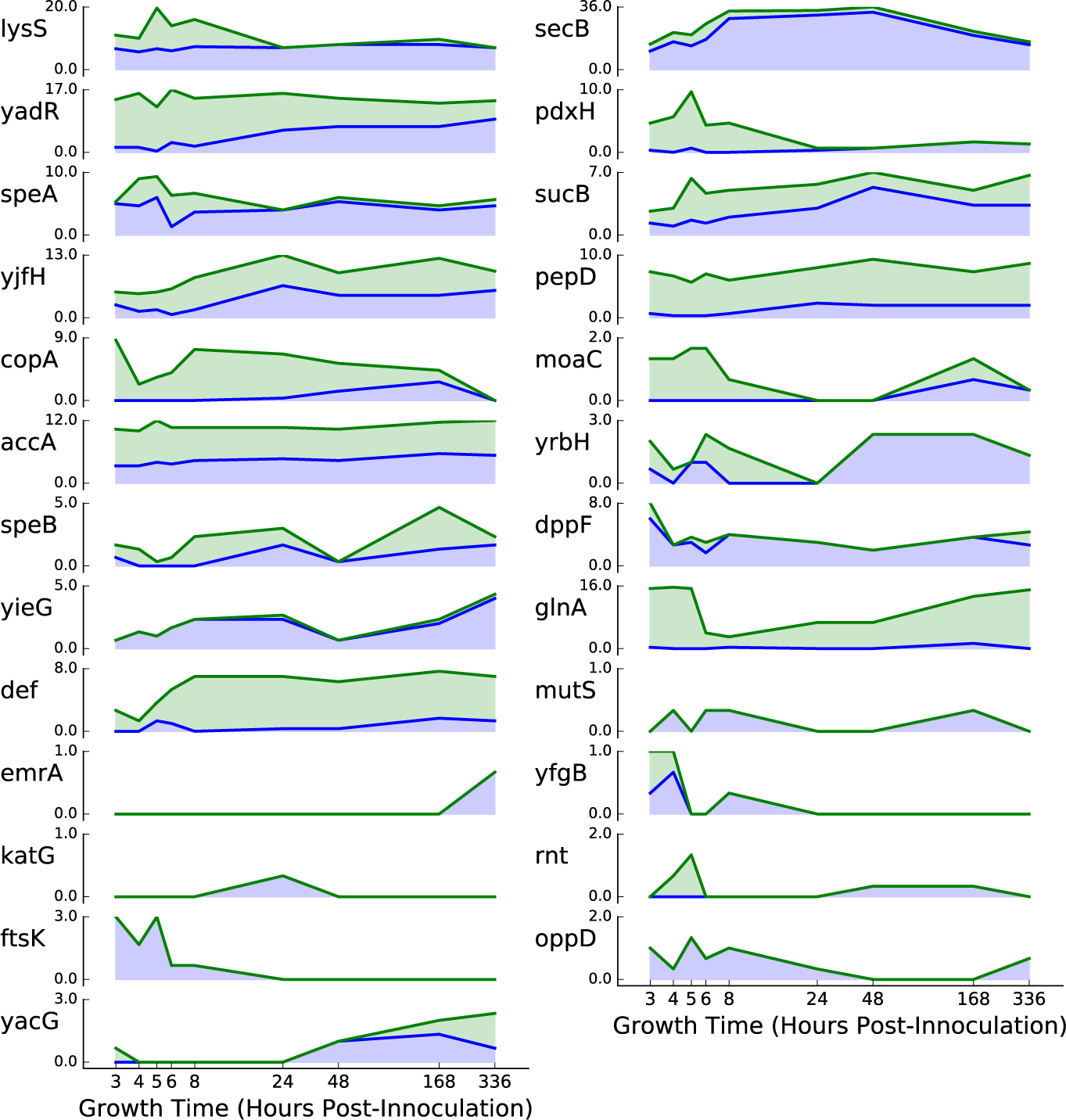
Temporally variable modification for individual proteins with an N-terminal serine possessing a +42 Da modification. Plots show unmodified (green) and +42 Da Modified (blue) PSM counts across all nine timepoints (*x*-axis) for the N-terminal position of all proteins that have both (i) at least one PSM identified by MODa as containing an N-terminal +42 Da modification and (ii) having a penultimate serine (AA position 2; i.e. the N-terminal residue following N-terminal methionine cleavage). Counts represent the average of the three biological replicates. Plots are ordered from top to bottom by the mean *p* value of the Fisher’s exact test for preferential modification (see text) from left-to-right within each row, and top-to-bottom across rows, with the most significant protein at the top left.

**Figure S5.**
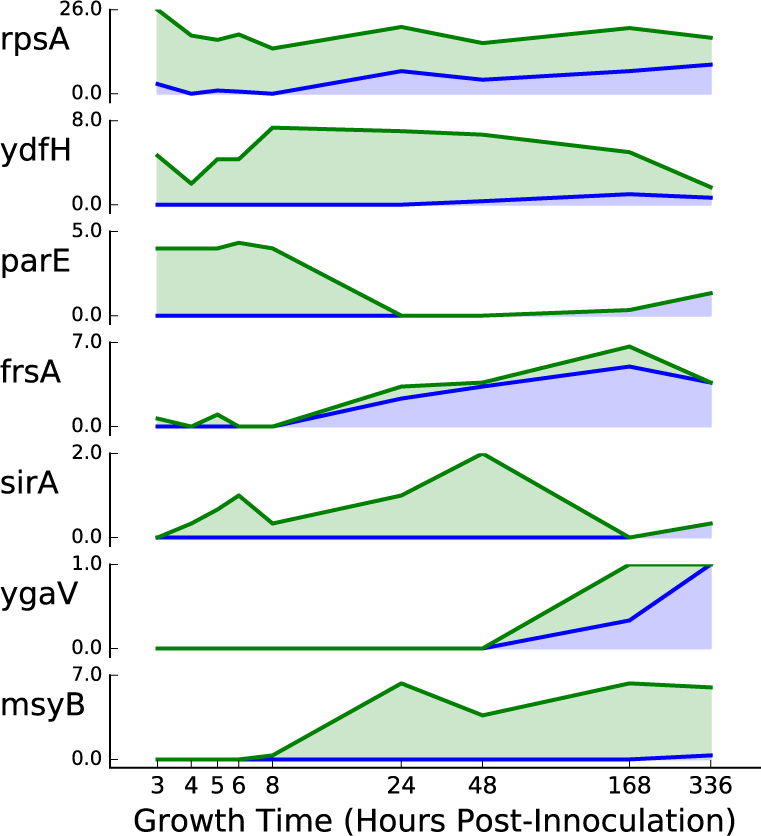
Temporally variable modification for individual proteins with an N-terminal threonine possessing a +42 Da modification. Plots show unmodified (green) and +42 Da Modified (blue) PSM counts across all nine timepoints (*x*-axis) for the N-terminal position of all proteins that have both (i) at least one PSM identified by MODa as containing an N-terminal +42 Da modification and (ii) having a penultimate threonine (AA position 2; i.e. the N-terminal residue following N-terminal methionine cleavage). Counts represent the average of the three biological replicates. Plots are ordered from top to bottom by the mean *p* value of the Fisher’s exact test for preferential modification (see text) from top to bottom, with the most significant protein at the top.

**Figure S6.**
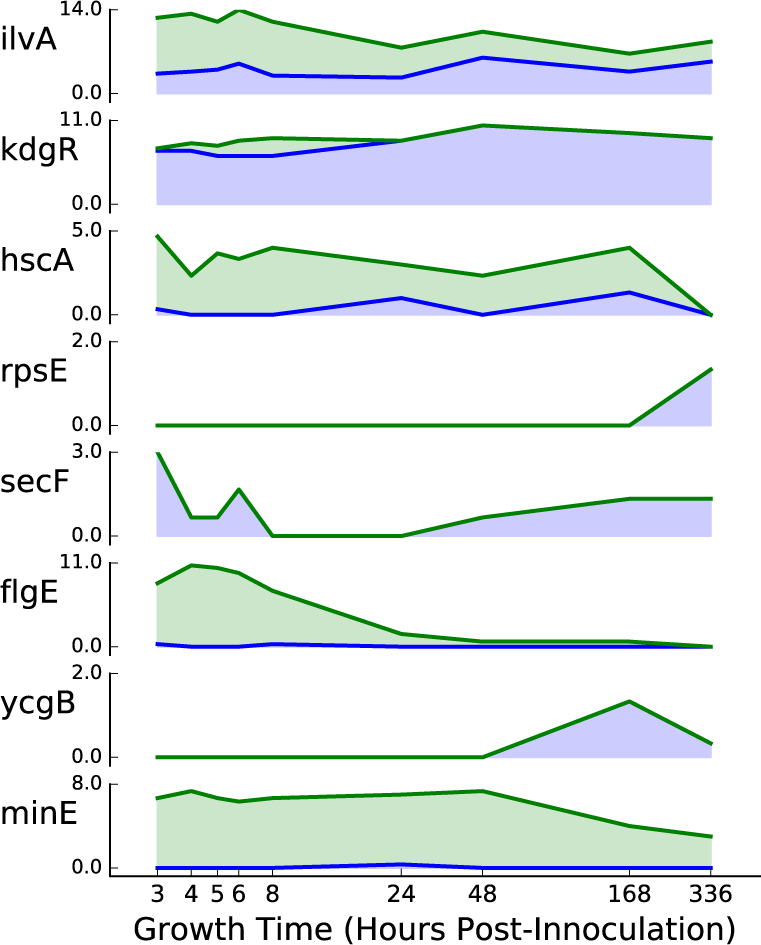
Fraction of total peptides across timepoints with an N-terminal alanine possessing a +42 Da modification. Plots show unmodified (green) and +42 Da modified (blue) PSM counts across all nine timepoints (*x*-axis) for the N-terminal position of all proteins that have both (i) at least one PSM identified by MODa as containing an N-terminal +42 Da modification and (ii) having a penultimate Alanine (AA position 2; i.e. the N-terminal residue following N-terminal methionine cleavage). Counts represent the average of the three biologcial replicates. Plots are ordered from top to bottom by the mean *p* value of the Fisher’s exact test for preferential modification (see text) from top to bottom, with the most significant protein at the top.

**Figure S7.**
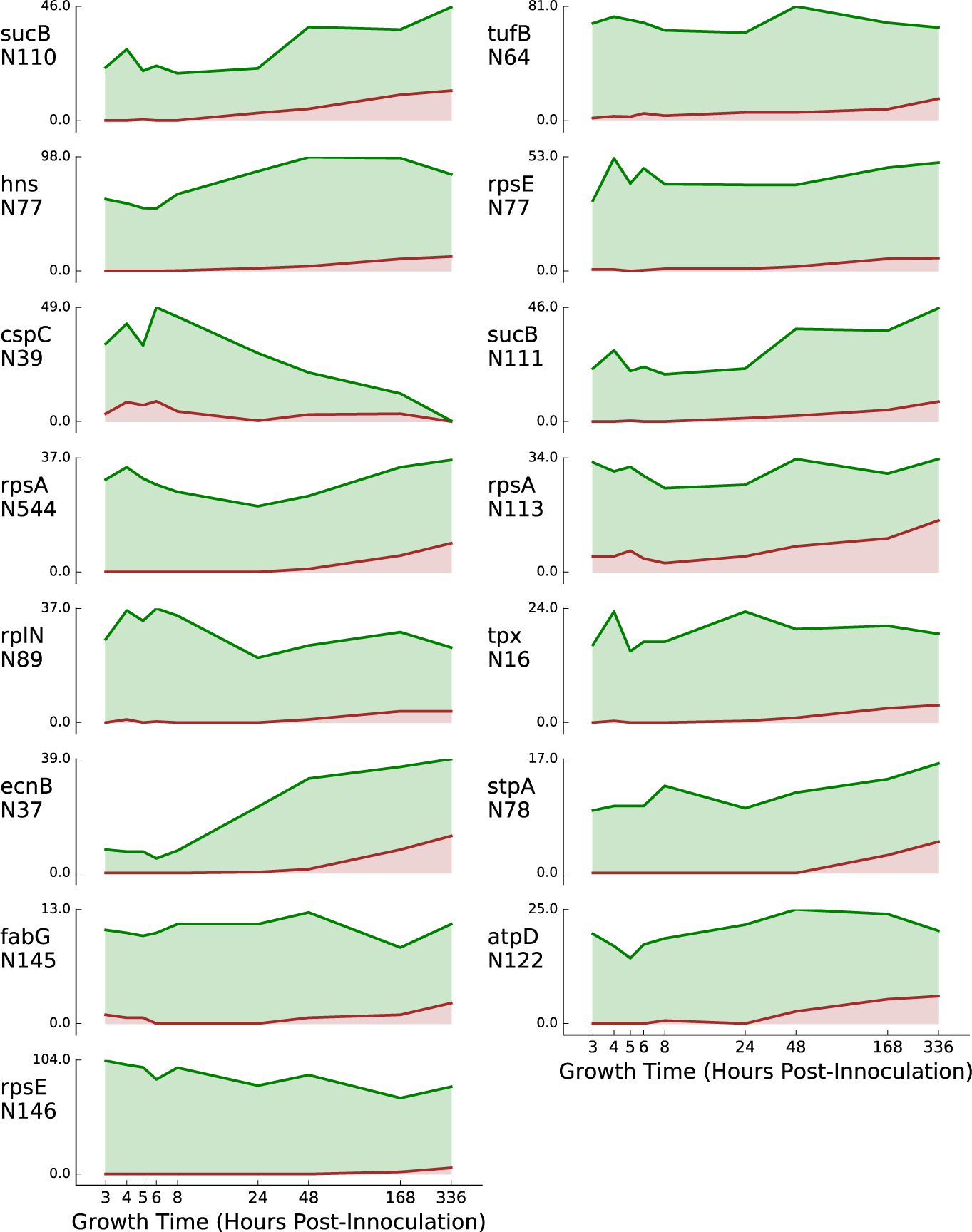
Modified and unmodified PSM counts for each AA position with a significantly stationary-phase biased +1 Da modification to asparagine. Plots show unmodified (green) and +1 Da modified (brown) PSM counts across all nine timepoints (*x*-axis) for the 10 asparagine residues with the most significant *p*-values across all three biological replicates. Counts represent the average of the three biologcial replicates. Plots are ordered by the mean *p* value of the Fisher’s exact test for preferential modification from left-to-right within each row, and from top-to-bottom across rows, with the most significant position at the top left.

**Figure S8.**
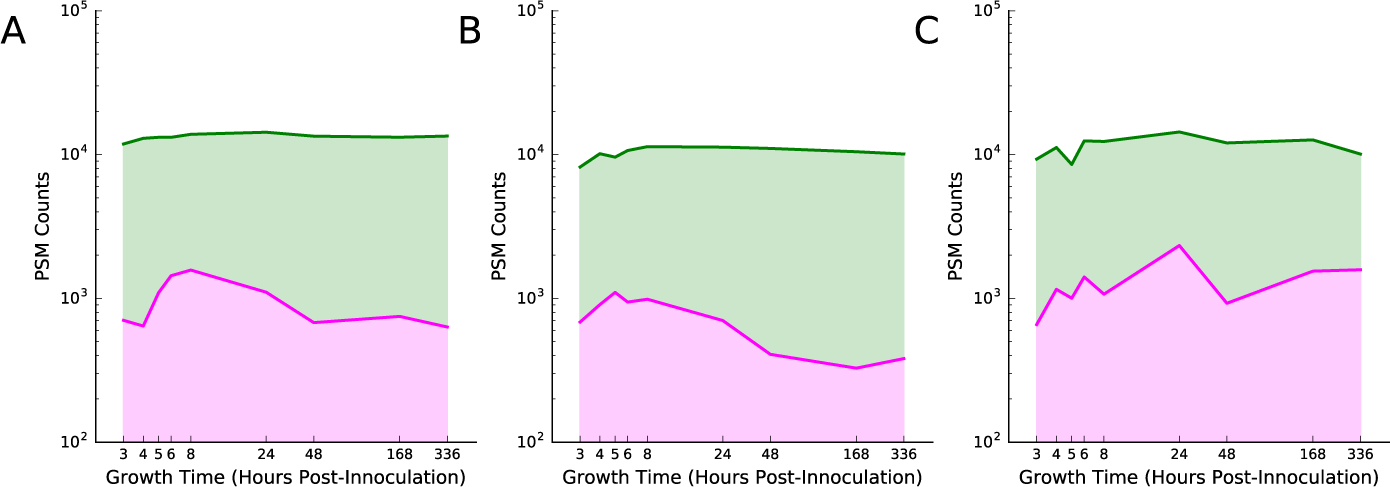
Modified and unmodified counts across timepoints for all AA positions with a +16 Da modification to methionine, pooled by biological replicate. Plots show the total unmodified (green) and +16 Da modified (magenta) PSM counts across all nine timepoints (*x*-axis) for methionine residues that have at least one +16 Da modification at any time point in any replicate. The three panels show counts for each of the three biological replicates (A), 2 (B) and 3 (C). Note that the *y*-axis is plotted on a logarithmic (base 10) scale due to the high number of total counts relative to modified counts.

**Figure S9.**
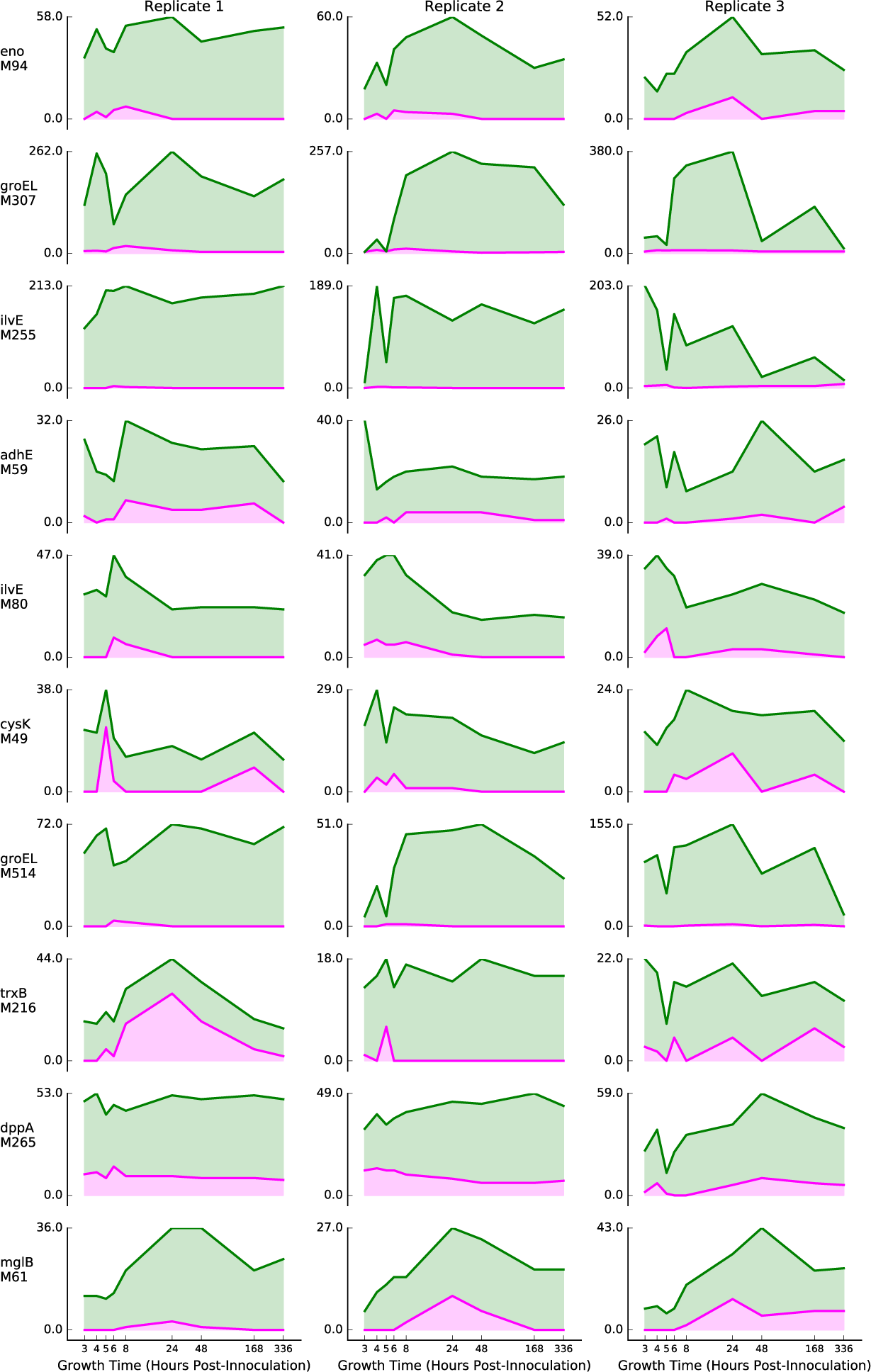
Modified and unmodified counts across timepoints for the top 10 exponential-enriched AA positions with a +16 Da modification to methionine. Plots show unmodified (green) and +16 Da modified (magenta) methionine PSM counts across all nine timepoints (*x*-axis) for the protein and position indicated. Plots in columns correspond to the three biological replicates 1 (left column), 2 (center column), and 3 (right column). Counts represent the average of the three biologcial replicates. Plots are ordered from top to bottom by the mean *p* value of the Fisher’s exact test for preferential modification (see text), with the most significant protein at the top.

**Figure S10.**
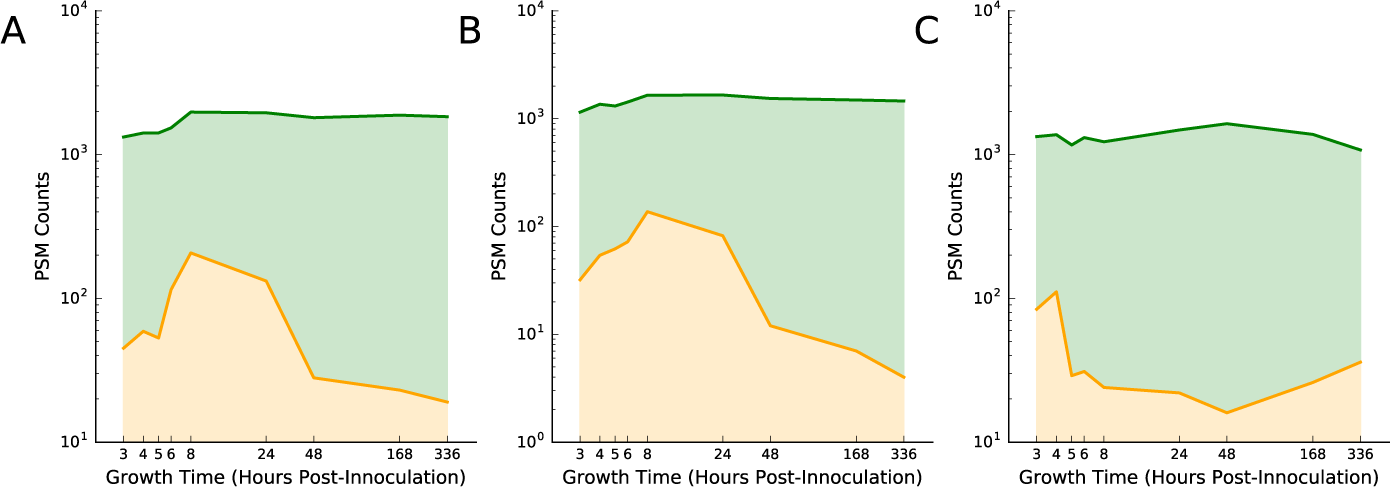
Modified and unmodified counts across timepoints for all AA positions with a +16 Da modification to tryptophan, pooled by biological replicate. Plots show the total unmodified (green) and +16 Da modified (orange) PSM counts across all nine timepoints (*x*-axis) for tryptophan residues that have at least one +16 Da modification at any time point in any replicate. The three panels show counts for each of the three biological replicates 1 (A), 2 (B) and 3 (C). Note that the *y*-axis is plotted on a logarithmic (base 10) scale due to the high number of total counts relative to modified counts.

**Figure S11.**
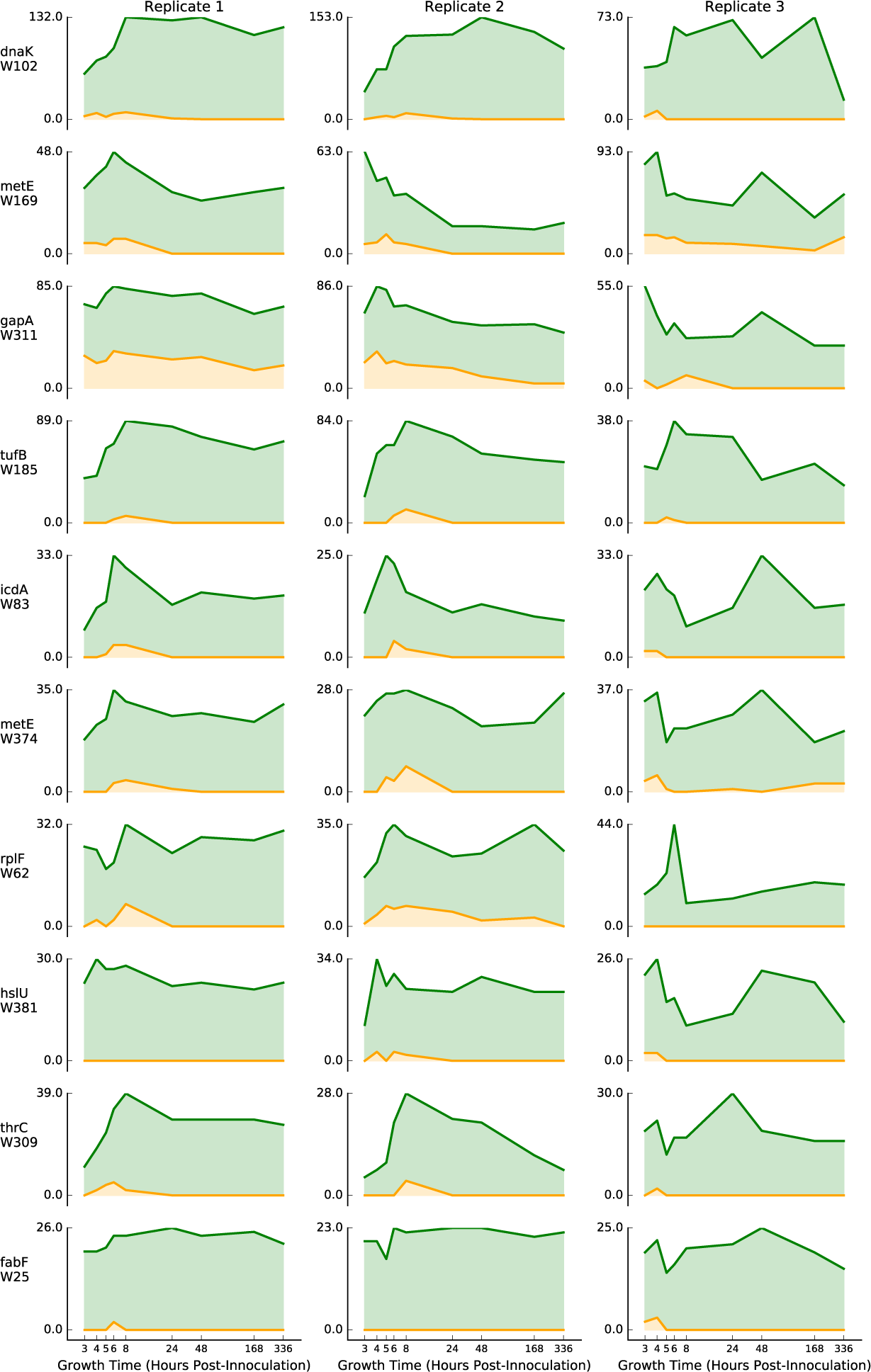
Modified and unmodified counts across timepoints for the top 15 exponential-enriched AA positions with a +16 Da modification to tryptophan. Plots show unmodified (green) and +16 Da modified (orange) tryptophan PSM counts across all nine timepoints (*x*-axis) for the protein and position indicated. Plots in columns correspond to the three biological replicates 1 (left column), 2 (center column), and 3 (right column). Counts represent the average of the three biologcial replicates. Plots are ordered from top to bottom by the mean *p* value of the Fisher’s exact test for preferential modification (see text), with the most significant protein at the top.

Additional file 12—Data Tables

Zip file containing several data tables in tab-separated format, as well as a readme file that explains the contents of each data file.

